# Physiological genomics of dietary adaptation in a marine herbivorous fish

**DOI:** 10.1101/457705

**Authors:** Joseph Heras, Mahul Chakraborty, J.J. Emerson, Donovan P. German

## Abstract

Adopting a new diet is a significant evolutionary change and can profoundly affect an animal’s physiology, biochemistry, ecology, and its genome. To study this evolutionary transition, we investigated the physiology and genomics of digestion of a derived herbivorous fish, the monkeyface prickleback (*Cebidichthys violaceus*). We sequenced and assembled its genome and digestive transcriptome and revealed the molecular changes related to important dietary enzymes, finding abundant evidence for adaptation at the molecular level. In this species, two gene families experienced expansion in copy number and adaptive amino acid substitutions. These families, amylase, and bile salt activated lipase, are involved digestion of carbohydrates and lipids, respectively. Both show elevated levels of gene expression and increased enzyme activity. Because carbohydrates are abundant in the prickleback’s diet and lipids are rare, these findings suggest that such dietary specialization involves both exploiting abundant resources and scavenging rare ones, especially essential nutrients, like essential fatty acids.

## Main

Populations exposed to new environments often experience strong natural selection (e.g., Herrel et al. 2008). Comparing closely related species has been an effective perspective in pinpointing changes that drive adaptation (Dasmahapatra et al. 2012; Lamichhaney et al. 2015). In its most powerful incarnation, the comparative method links variation at the genetic level to molecular phenotypes that in turn change how whole organisms interact with their environments, revealing for example adaptation to new abiotic factors (Protas et al. 2006; Chakraborty and Fry 2015; Peichel and Marques 2017; Tong et al. 2017; Chen et al. 2018), changes in diet (Harris and Munshi-South 2017; Hsieh et al. 2017; Zepeda-Mendoza et al. 2018), and exposure to pollution (Vega-Retter et al. 2018). In animals, digestion is an ideal model phenotype because it is central to fitness, is understood in many species at genetic, molecular, biochemical, and physiological levels, and is a trait that shows abundant variation throughout animal evolution (Karasov and Martinez del Rio 2007). While studies of animal digestion have yielded insights into the physiology and biochemistry of adaptation, untangling its genetic basis is contingent on the availability and quality of genomic resources, which have traditionally been lacking in nonmodel species. Advances in genome technology have improved the quality and decreased the cost of obtaining sequences, stimulating the production of genome assemblies and catalyzing genetic discoveries for a diverse array of non-model organisms.

Here we describe the genetic changes accompanying acquisition of an herbivorous diet in the marine intertidal fish, *Cebidichthys violaceus*. To do this we generated a physiological genomics dataset for this non-model species, including a highly contiguous and complete genome, the transcriptomes of digestive and hepatic tissues, and digestive enzyme activity levels. We generated a reference quality genome for *C. violaceus* with an N50 of 6.7 MB, placing it among the most contiguous teleost assemblies (Fig. 1). The ecological and evolutionary positions of *C. violaceus* makes these resources particularly suited to unraveling the acquisition of herbivory. Members of the family Stichaeidae, including *C. violaceus*, have independently invaded the intertidal zone multiple times, and in two cases, began consuming significant amounts of algae (Fig. 2; Kim et al. 2014), a diet that is low in protein and lipids and rich in fibrous cell walls (Horn et al. 1986; German et al. 2015). In addition to this convergent acquisition of herbivory, these intertidal invaders experience extremes of environment, including osmotic variability, wide temperature fluctuations, and aerial exposure.

**Fig. 1.**
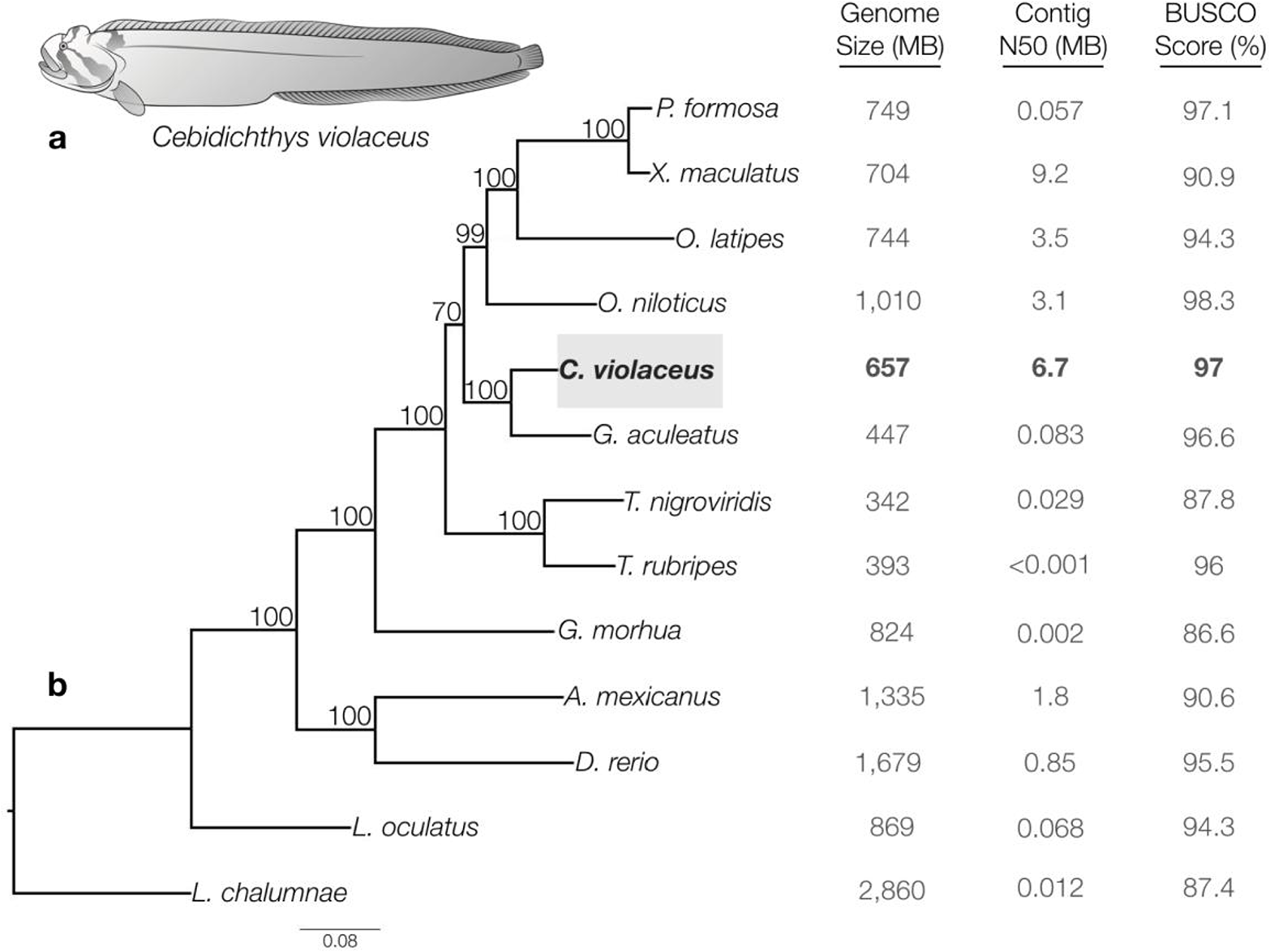
Phylogenetic relationships from 13 fishes with sequenced genomes including the monkeyface prickleback, *Cebidichthys violaceus*. **a**, Illustration of the monkeyface prickleback (*Cebidichthys violaceus*). ***b***, A maximum likelihood (ML) tree was constructed with 1,000 bootstrap replicates in PhyML v3.1 based on the lowest average gaps present in our ortholog cluster alignments of concatenated 30 protein coding genes from all 13 taxa. *C. violaceus* (highlighted in a gray box) along with 12 fish taxa with sequenced genomes from Ensembl (Release 91) were used for our phylogenetic analyses. The Ensembl taxa included: *Poecilia formosa*, *Xiphophorus maculatus*, *Oryzias latipes*, *Oreochromis niloticus*, *Gasterosteus aculeatus*, *Tetraodon nigroviridis*, *Takifugu rubripes*, *Gadus morhua*, *Astyanax mexicanus*, *Danio rerio*, *Lepisosteus oculatus*, and *Latimeria chalumnae*. Genome size and contig N50, and BUSCO v3 complete genes identified out of 2586 BUSCO groups are represented for all taxa.

**Fig. 2.**
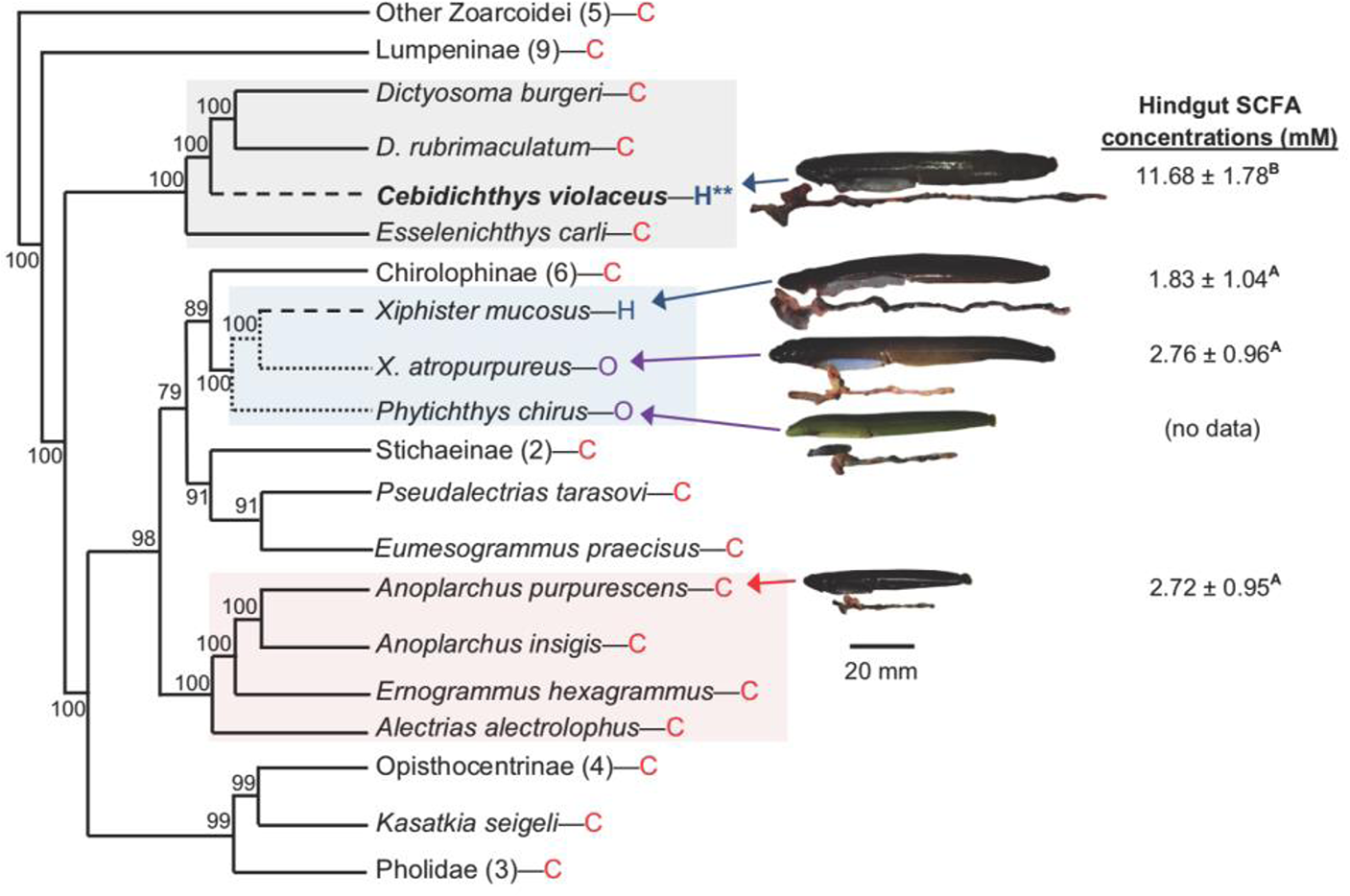
Phylogenetic relationships of the polyphyletic family Stichaeidae based on 2,100 bp of *cytb, 16s*, and *tomo4c4* genes (Kim et al. 2014). Bayesian posterior probabilities are indicated on nodes. *Cebidichthys violaceus* is bolded, and photos of *C. violaceus* and other studied taxa are shown with their digestive systems beneath their bodies. Note the differences in gut size. H=herbivory, O=omnivory, C=carnivory. Evolution of herbivory (— — — —) and omnivory (…………) are shown. Numbers in parentheses show number of taxa evaluated at that branch. Boxes highlight alleged families or subfamilies within the polyphyletic family Stichaeidae, with Cebidichthyidae (top), Xiphisterinae (middle), and Alectriinae (bottom) all highlighted. Hindgut short chain fatty acid (SCFA) concentrations are mean ± standard deviation, and were compared with ANOVA (F_3,33_ = 127.92; *P* < 0.001). SCFA data from German et al. (2015).

Herbivory is poorly represented among high quality teleost genomes (Fig. 1; Supplementary Table S1). Because teleosts are so speciose, they represent a large number of independent acquisitions of herbivory, even though only 5% of teleosts are considered nominally herbivorous (cf 25% for mammals; Choat and Clements 1998). Even the term “herbivorous” is controversial amongst ichthyologists (Clements et al. 2017). Among nominally herbivorous fishes, most do not specialize on algal thalli like *C. violaceus* does. Moreover, just within the Stichaeidae, it is clear that *C. violaceus* digests red and green algae with the aid of microbial symbionts in their hindguts, as evidenced by elevated levels of short chain fatty acids, or SCFAs, in this gut region. This microbial symbiosis is analogous to other highly specialized vertebrate herbivores like lagomorphs or rodents, whereas the other stichaeid herbivore, *X. mucosus*, is less- reliant on such microbes (Fig. 2; German et al. 2015). Thus *C. violaceus* offers a unique opportunity to study extremes of dietary specialization. Although herbivorous and omnivorous animals tend to have elevated amylase activities in their guts (Perry et al. 2007; German et al. 2010; Kohl et al. 2011; Axelsson et al. 2013; Boehlke et al. 2015; German et al. 2015), and achieve these activities via gene duplications or elevated expression of fewer amylase genes (German et al. 2016), the appreciation of bile-salt activated lipase in the digestive process of animals eating a low-lipid, high-fiber food, is more recent (German et al. 2004; German et al. 2015; Leigh et al. 2018). Here we describe extensive structural variation at the gene level and adaptive variation at the amino acid level in both of these gene families, suggesting multiple mutational mechanisms for the acquisition of a novel derived dietary physiology in *C. violaceus*.

## Results and Discussion

### Genome Assembly, Quality, and Size

We generated ~30Gb long reads (~37X based on genome size = 792 Mb for *C. violaceus)* (Hinegardner and Rosen, 1972) using Pacific Biosciences (PacBio) Single Molecule Real Time sequencing, and 84.5Gb (~107X) paired end Illumina reads. A draft assembly using the Illumina data alone was highly fragmented (N50 = 2760 bp), consistent with most published fish genome assemblies [Zerbino et al. 2017; www.ensembl.org]. Using the Illumina contigs in concert with long reads yielded a markedly more contiguous hybrid assembly (N50 =2.21 Mb). An assembly of the long reads alone yielded a similarly contiguous genome (N50 = 2.45 Mb). Finally, merging of the hybrid assembly with the long read only assembly (Chakraborty et al. 2016; see Methods) yielded a highly contiguous assembly (N50 = 6.69Mb), ranking it among the most contiguous teleost genomes, and, to our knowledge, the most contiguous among herbivorous fishes. Assessment of the universal single copy orthologs (BUSCOs) (Waterhouse et al. 2017) show that completeness of our assembly (97%) is comparable to or even better than the model reference fish genomes (86.6-98.3 %) (Fig. 1B). By using JELLYFISH we estimated the genome size, based on an average of four k-mer size counts (25, 27, 29, 31), at 656,598,967 base pairs based with a standard deviation of 4,138,853 base pairs (Supplementary Figure S5; Supplementary Table S4), which is close to the original genome size estimate of 792 Mb based on the c-value (Hinegardner and Rosen, 1972), and for other fish genomes (Fig. 1B; Zerbino et al. 2017).

Consistent with phylogenetic considerations, the genome of *C. violaceus* shares the most similarity with that of stickleback (Supplementary Figure S16). When we compare our assembled genome to those of *Lepisosteus oculatus*, *Danio rerio*, and *O. latipes*, large sections of syntenic regions are conserved (Supplementary Figure S16-19). Nevertheless, these comparisons show the relative completeness of our draft genome in comparison to these model systems.

### Physiological genomics of digestive enzymes

The activity levels of digestive enzymes reveal what substrates are readily digested in an animal’s digestive tract, highlighting the enzyme genes that are potential targets of selection for efficient digestion (Karasov and Martínez del Rio 2007). Within the field of Nutritional Physiology, the Adaptive Modulation Hypothesis predicts a match between the amount of an ingested substrate (e.g., starch) and digestive enzyme expression and activity levels to digest such substrates (e.g., amylase) based on economic principles (Karasov and Martinez del Rio 2007). Indeed, pancreatic amylase activity tends to be elevated in the guts of herbivores and omnivores in comparison to carnivores (especially in prickleback fishes; Chan et al. 2004; German et al. 2016), matching the higher intake, and the importance, of soluble carbohydrates to these animals (Horn et al. 2006; Kohl et al. 2011; German et al. 2010; Axelsson et al. 2013; Boehlke et al. 2015; German et al. 2015; German et al. 2016). Our assembly reveals three tandem pancreatic amylase genes in the *C. violaceus* genome: two copies of *amy2a* and one copy of *amy2b* (Fig. 4). The three *amy* genes in tandem differs from other pricklebacks (and indeed, most other fishes for which genetic data are available), which tend to have one or two identical copies of *amy2* (Fig. 4; German et al. 2016). The two *amy2a* copies are supported by three spanning reads, emphasizing the correct assembly of the *amy2a* tandem duplicates (Supplementary Figure S12). Each amylase gene is preceded by a 4.3Kb DNA element encoding a transposase (Fig. 4, Supplementary Figure S13), hinting at a role of this TE in gene duplications in this region (Feschotte and Pritham 2007; McVean 2010; Pantzartzi et al. 2018). Additionally, the *amy2B* gene has a 2,025 bp LINE element inside the 2nd intron of the gene. All three copies of amylase gene copies possess a ~470bp fragment of a long terminal repeat retrotransposon (Supplementary Figure S13). Insertion of the TEs proximal to the transcription start site and within first introns could modulate the expression of the amylase gene copies because both of these regions are typically enriched with cis-regulatory elements [Feschotte, 2008]. When testing all 11 branches for the seven prickleback taxa, only one branch was under episodic diversifying selection (C. *violaceus, amy2b*) with a significant uncorrected p-value of 0.0044 (Figure 4B). We do not see this pattern of positive selection in any of the other branches with significant p-values. We observed three sites with episodic positive selection with a p-value under 0.05 (sites: 41, 256, and 279; Figure 4D). How the AMY2A and AMY2B proteins differ in function requires further investigation, but they have different isoelectric points (7.86 vs. 8.62; German et al. 2016), which hints that they may be more active in different parts of the gut, and indeed, the transcriptomic data show that *amy2a* is expressed at a fairly constant level throughout the proximal GI tract (including pancreatic tissue in the pyloric ceca), whereas *amy2b* is expressed mostly in the mid intestine (Fig. 3).

**Fig. 3.**
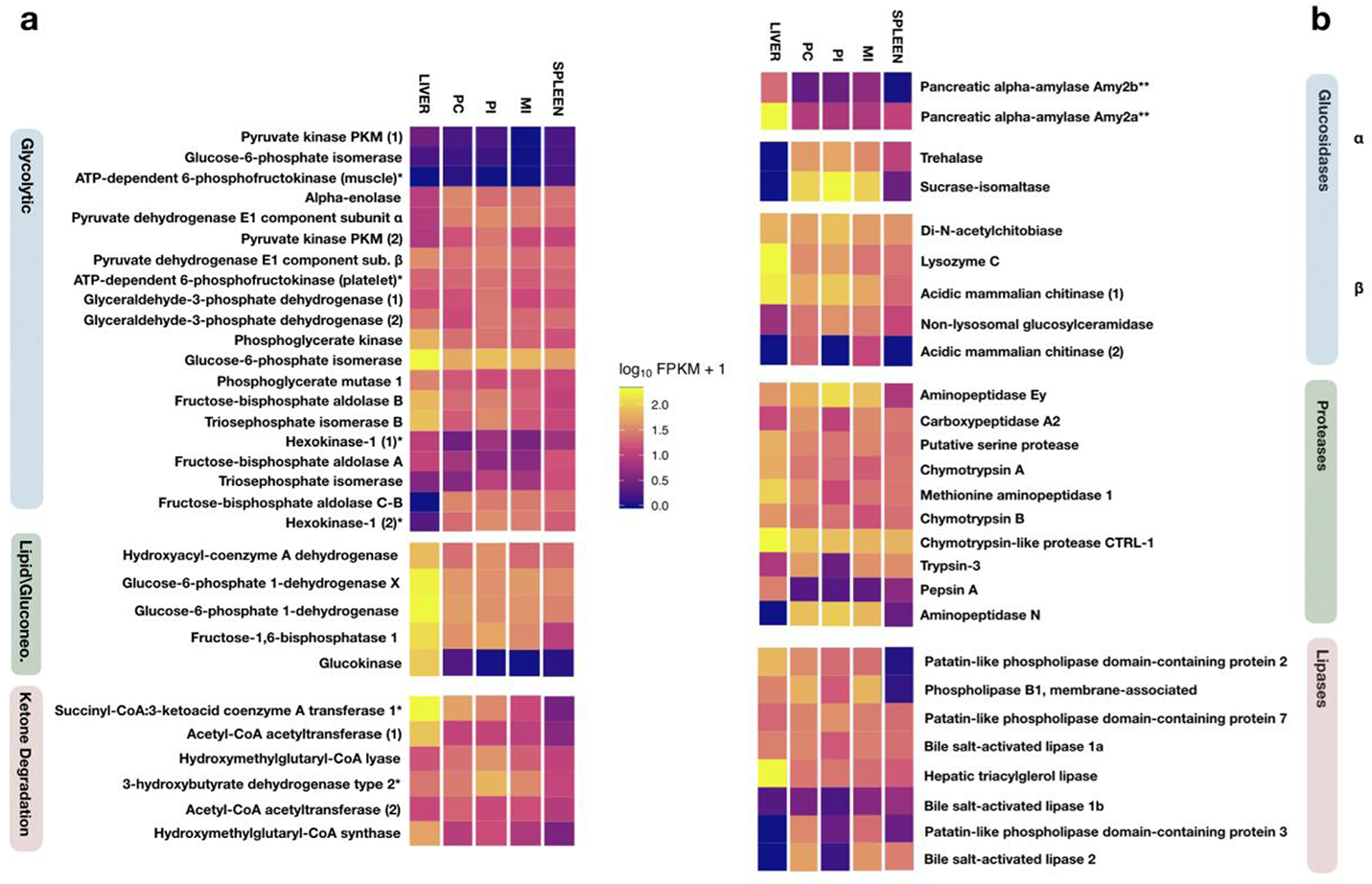
Gene expression profiles of nine tissue types from the *Cebidichthys violaceus* transcriptome. We used brain, gill, gonads (testes), heart, liver, pyloric caeca (PC), proximal intestine (PI), and middle intestine (MI), and spleen tissues from *C. violaceus* to represent the transcriptome. Only gene expression profiles of liver, PC, PI, MI and spleen are represented for Glycolytic, Lipid metabolism/Gluconeogenesis, Ketone Degradation, Glucosidases, Proteases, and Lipases. Low to high expression is shown on a gradient scale from violet to yellow respectively. Unit of expression is measured as Fragments Per Kilobase of transcript per Million mapped reads (FPKM). **a**, Three heatmaps were generated for transcripts which belong to glycolytic pathways (blue box), transcripts for enzymes associated with gastrointestinal fermentation based on Willmott et al. 2005 study in marine teleost fishes (green box), and kentone degradation pathway transcripts (red/pink box). **b**, Three heatmaps of digestive enzymes were generated which include carboyxlases (alpha-glucosidases and beta glucosidases, blue boxes), proteases (green box), and lipases (red/pink box).

**Fig. 4.**
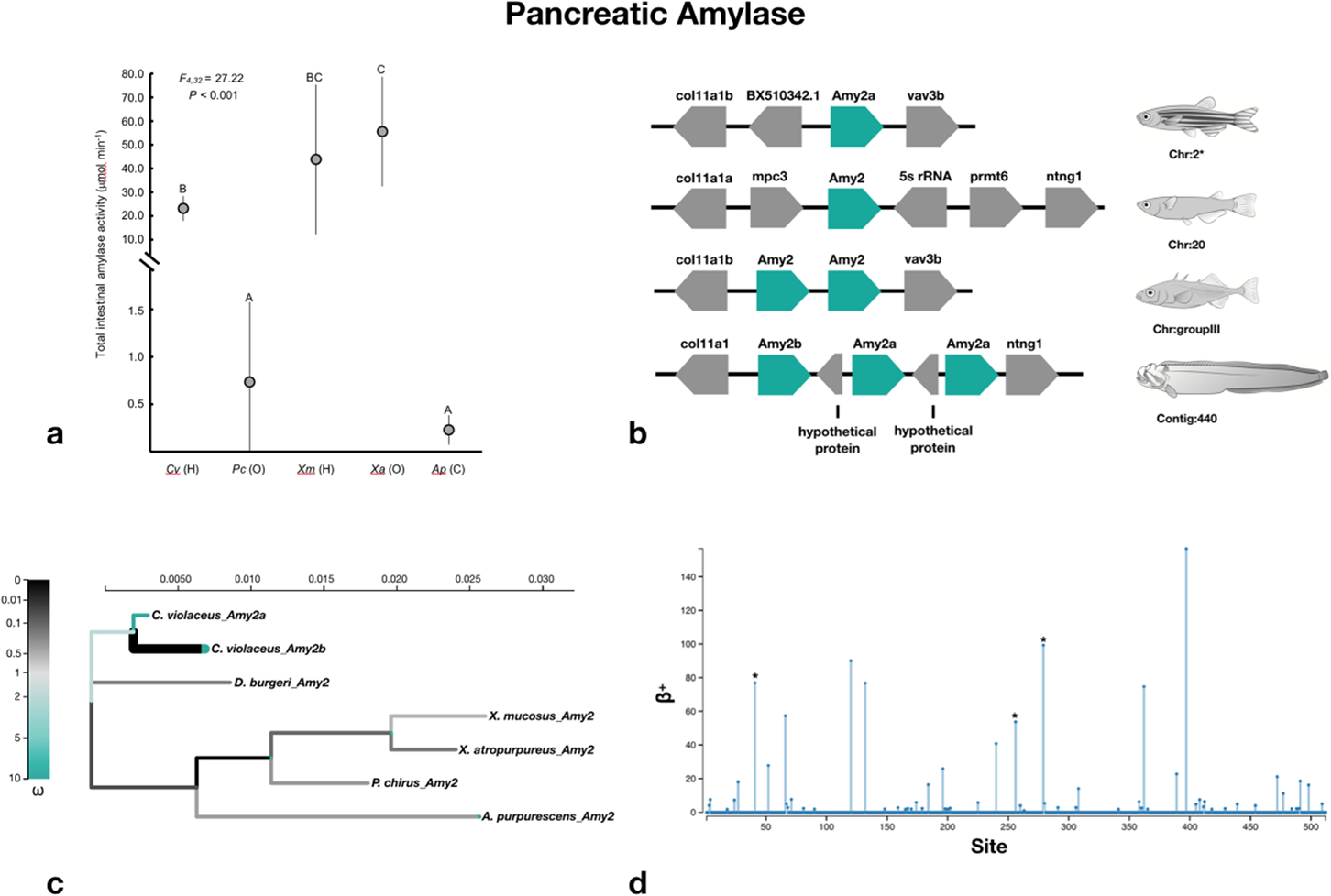
Enzyme activity, gene copy number, and molecular evolution of amylase (a carbohydrase). **a**, Total gut standardize enzymatic activity for *Cebidichthys violaceus* (Cv) and other intertidal stichaeid species: *Phyticus chirus* (Pc), *Xiphister mucosus* (Xm), *Xiphister atropurureus* (Xa), and *Anoplarchus purpurescens* (Ap). H = herbivory, O = Omnivory, and C = Carnivory. Values are mean ± standard error with n = 6 for Cv, Xm, Xa, and Ap; and n = 9 for Pc (German et al. 2015). Interspecific comparisons were made for amylase with ANOVA, where circles that share a letter are not significant. **b**, Synteny map for amylase genes from *Danio rerio, Oryzias latipes, Gasterosteus aculeatus*, and *C. violaceus*. * denotes that *D. rerio* also has amylase loci present on chromosome 17. **c**, An adaptive branch-site Random Effects Likelihood (aBSREL) test for episodic diversification phylogenetic tree constructed for amylase genes from *C. violaceus* (amy2a and amy2b) and other intertidal stichaeid species. ω is the ratio of nonsynonymous to synonymous substitutions. The color gradient represents the magnitude of the corresponding ω. Branches thicker than the other branches have a P<0.05 (corrected for multiple testing) to reject the null hypothesis of all ω on that branch (neutral or negative selection only). A thick branch is considered to have experienced diversifying positive selection. **d**, The output of Mixed Effects Model of Evolution (MEME) to detect episodic positive/diversifying selection at sites. β+ is the non-synonymous substitution rate at a site for the positive/neutral evolution throughout the sequence of the gene. ** is an indication that the positive/diversifying site is statistically significant with a p-value < 0.01 and * is for p-value < 0.05.

Interestingly, amylolytic activities in the guts of *C. violaceus* are similar to those in the two species of *Xiphister* (Figs. 2 and 4a), yet the *Xiphister* taxa only have two copies of *amy2a* (German et al. 2016), and *C. violaceus* and *X. mucosus* digest algal starch with similar efficiencies (Horn et al. 1986). Thus, the phenotype of elevated amylase activity can be achieved via different mechanisms (increased gene copy number leading to increased expression of the genes vs increased expression of fewer genes) with similar performance outcomes at the whole animal level (Horn et al. 1986; German et al. 2016).

Herbivores have the challenge of consuming a food that is simultaneously low in lipid (Neighbors and Horn 1991), and high in fiber (Painter 1983), and fiber binds to lipid, impeding its digestion (German et al. 1996). Thus, Bile Salt Activated Lipase (BSAL; Murray et al. 2003) represents another important digestive enzyme for herbivorous fishes because lipids (especially essential fatty acids) are essential to survival. The importance of BSAL in herbivores has been validated in prickleback fishes (German et al. 2004; German et al. 2015), as well as in *Danio rerio* fed a high-fiber, low-lipid diet analogous to an herbivorous diet in the laboratory, which elicited elevated lipase activities in this species (Leigh et al. 2018). Thus, it appears that *C. violaceus*, the algal-consuming *Xiphister* taxa, and other herbivorous species (German et al. 2004; Leigh et al. 2018) invest in lipase expression to ensure lipid digestion from their algal diet, consistent with what is known as the Nutrient Balancing Hypothesis, or that animals can invest in the synthesis of digestive enzymes to acquire limiting nutrients (Clissold et al. 2010), in this case, lipids.

In the *C. violaceus* genome, we identified four tandem copies of Bile Salt Activated Lipase (BSAL) on contig 445 (Figure 5B). Two of these (BSAL-1b and BSAL-1c) are more similar to each in intron/exon arrangements as compared to BSAL-1a and BSAL-2, suggesting that BSAL-1b and BSAL-1c copies originated from more recent gene duplications (Supplementary Figure S14). Interestingly, BSAL-1a and BSAL-2 possess two different LINE elements in their first and last introns, respectively, contributing to the structural diversification of the two gene copies (Supplementary Figure S14). In addition, there is evidence of recombination in BSAL, and there is one breakpoint (with a p-value of 0.01) with significant topological incongruence at site 534. To examine whether BSAL-2 diverged under selection, we estimated selection with 11 taxa and found three branches out of 19 to be under episodic diversifying selection: *C. violaceus* BSAL2 (uncorrected p-value 0.0008), Node 9 (uncorrected p-value 0.00001; which leads to the clade containing BSAL genes of *Phytichthys chirus* and the two species of *Xiphister*), and *X. mucosus* BSAL2 (uncorrected p-value 0.00001) (Fig. 5C). Interestingly, these all lead to taxa that consume algae, and hence, a lower lipid diet. In the lipase genes, there are 14 sites under episodic positive selection (p-value < 0.05): site 92, 94, 115, 139, 144, 151, 156, 160, 356, 429, 436, 443, 460, and 475 (Figure 5D). Consistent with their functional divergence in protein sequence, BSAL-1 and BSAL-2 show different tissue-specific expression patterns: BSAL-1a is expressed ubiquitously, whereas BSAL-2 is primarily expressed in pyloric caeca, spleen, and intestine [Fig. 3B]. Furthermore, even though the protein sequence of BSAL-1 copies are similar (pairwise distance of 0.014-0.054; poisson corrected), BSAL-1b is expressed largely in the gills and the heart, providing evidence for subfunctionalization in the BSAL-1 gene copies. Whereas *C. violaceus* has four copies of BSAL in their genome, other fishes for which the genome has been sequenced, and other stichaeids (based on transcriptomes of their relevant tissues), appear to have two BSAL genes (Fig. 5). These extra lipase genes may have real consequences for *in vivo* lipid digestibility, as *C. violaceus* digests algal lipid with consistent efficiency across a range of lipid concentrations, whereas *X. mucosus* shows decreasing lipid digestibility with decreasing lipid content in their diet (Horn et al. 1986). Humans who evolutionarily inhabited high latitudes in the Northern Hemisphere are known to consume lipid-rich diets, and genomic scans of some of these populations (Nganasans and Yakuts) shows that they have experienced selection on lipases and proteins involved in lipid metabolism (Hsieh et al. 2017). Much like amylase activities being elevated in herbivores and omnivores, selection on lipid digestion and metabolism in animals (in this case, humans) consuming high-lipid diets concurs with the Adaptive Modulation Hypothesis (Karasov and Martinez del Rio 2007), which is the opposite of the Nutrient Balancing Hypothesis (Clissold et al. 2010) that we propose for selection on lipase in *C. violaceus*. Thus, specific pathways can be selected upon for different reasons: some to ensure adequate digestion and metabolism of an abundant resource (Adaptive Modulation Hypothesis), the other for the acquisition of a limiting one (Nutrient Balancing Hypothesis).

**Fig. 5.**
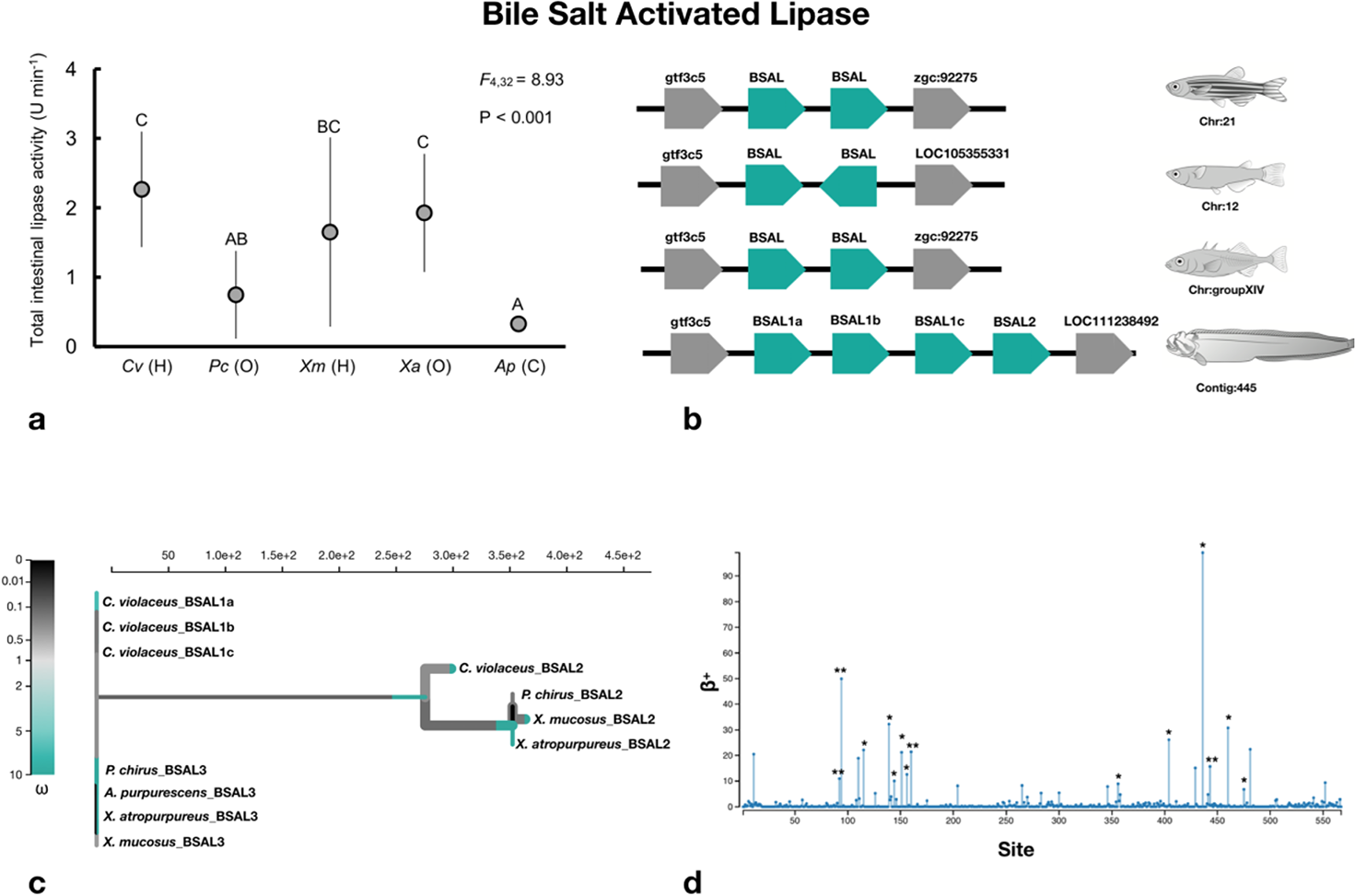
Enzyme activity, gene copy number, and molecular evolution of bile salt activated lipase (BSAL). **a**, Total gut standardize enzymatic activity for Cebidichthys violaceus (Cv) and other intertidal stichaeid species: Phyticus chirus (Pc), Xiphister mucosus (Xm), Xiphister atropurureus (Xa), and Anoplarchus purpurescens (Ap). H = herbivory, O = Omnivory, and C = Carnivory. Values are mean ± standard error with n = 6 for Cv, Xm, Xa, and Ap; and n = 9 for Pc (German et al. 2015). Interspecific comparisons were made for BSAL with ANOVA, where circles that share a letter are not significant. **b**, Synteny map for BSAL genes from Danio rerio, Oryzias latipes, Gasterosteus aculeatus, and C. violaceus. **c**, An adaptive branch-site Random Effects Likelihood (aBSREL) test for episodic diversification phylogenetic tree constructed for BSAL genes from C. violaceus and other intertidal stichaeid species. ω is the ratio of nonsynonymous to synonymous substitutions. The color gradient represents the magnitude of the corresponding ω. Branches thicker than the other branches have a P<0.05 (corrected for multiple testing) to reject the null hypothesis of all ω on that branch (neutral or negative selection only). A thick branch is considered to have experienced diversifying positive selection. **d**, The output of Mixed Efffects Model of Evolution (MEME) to detect episodic positive/diversifying selection at sites. β+ is the nonsynonymous substitution rate at a site for the positive/neutral evolution throughout the sequence of the gene. ** is an indication that the positive/diversifying site is statistically significant with a p-value < 0.01 and * is for p-value < 0.05.

### Transcriptomics

We constructed a transcriptomics dataset for nine tissues (Supplemental Figures S8-S11) in *C. violaceus*, focusing here on the liver, pyloric ceca (which includes pancreatic tissue; Kim et al. 2014; German et al. 2016), mid intestine, and spleen (Fig. 3). In these four tissues, we identified genes associated with metabolism (glycolysis, ketone metabolism, and fermentation) (Fig. 3A), and digestion (Fig. 3B). Like herbivorous mammals, *C. violaceus* has an active microbial community in their hindgut that ferments dietary substrates to short chain fatty acids (SCFAs)(Fig. 2; German et al. 2015). These SCFA are then absorbed by the host and used to generate ATP in various tissues (Bergman 1990; Karasov and Martinez del Rio 2007). Because SCFAs are ketones, animals reliant on microbial fermentation in their hindguts also have active ketotic pathways in their tissues (Karasov and Martinez del Rio 2007; Willmott et al. 2005). Indeed, genes coding for proteins that are part of ketone synthesis and degradation are upregulated in most tissues in *C. violaceus*, but especially in the liver (Fig. 3A), consistent with the digestive strategy utilizing microbial symbionts in this herbivorous fish (German et al. 2015). In addition to amylase and lipase (discussed above), there are clear expression patterns for carbohydrases, proteases, and lipases, confirming the suite of enzymes necessary to digest a range of nutrients (Fig. 3B). Interestingly, *C. violaceus* appears to express three separate chymotrypsin genes and only one trypsin gene. This may be consistent with their consumption of algal material, as carnivores (e.g., salmon) may invest more in trypsin expression (Rungruangsak-Torrissen et al. 2006), whereas herbivores (e.g., grass carp) may express more chymotrypsin (Gioda et al. 2017).

Interestingly, we found that the pyloric cecal tissue of *C. violaceus*, which is frequently recognized as “pancreatic” (e.g., German et al. 2016) because it is sheathed in acinar cells (Kim et al. 2014) and shows elevated activity levels of pancreatic enzymes (German et al. 2015), only had two differentially expressed genes in comparison to the mid intestine (Supplementary Figures S9-S11), which is known to be the highly absorptive region of the fish gut (Buddington et al. 1987; Tengjaroenkul et al. 2000). The pyloric ceca have been documented to be absorptive (Buddington and Diamond 1987), meaning they can be functionally similar to the mid intestine, but the mid intestine is rarely recognized as also having pancreatic function. The pancreatic tissue of fish does not generally form a distinct organ, as it does in mammals. In fishes, it tends to be embedded in the liver (forming a hepatopancreas) or diffuse along the intestine (Clements and Raubenheimer 2006). Our transcriptomic data show not only a shared absorptive function of the pyloric caeca, but also that the acinar cells are distributed along the intestine and not restricted to the pyloric cecal region (Supplementary Figures S10-S11). Indeed, the mid intestine shows expression of numerous pancreatic enzymes, the activities of which are detectable in mid intestine tissue of prickleback fishes (German et al. 2015). Thus, our transcriptomic and biochemical data strongly suggest that the pyloric ceca are absorptive (as suggested by Buddington and Diamond 1987) and that pancreatic tissue is diffuse along the intestine in prickleback fishes.

### Conclusions

Our genomic efforts produced a highly contiguous fish genome, and because of the rich history of investigation into the nutritional physiology of *C. violaceus* and other stichaeid fishes, we were able to analyze our genomic data in the context of digestive physiology. Our results are unique in that we show that both, the Adaptive Modulation Hypothesis (Karasov and Martinez del Rio 2007), and the Nutrient Balancing Hypothesis (Clissold et al. 2010), can have genetic underpinnings within the same organism. This powerful physiological genomics approach will allow us to tease apart more aspects of dietary adaptation in *C. violaceus* moving forward and to provide a model for nutritional physiological research. Given that *C. violaceus* is an important member of Marine Protected Areas on the west coast of the United States, and is targeted for aquaculture in northern California where it is a delicacy, our data will also have application for conservation and better culturing techniques.

## METHODS

### Collection and Preparation

One individual of *Cebidichthys violaceus* with a standard length of 156mm was collected in May 2015 by hand during low tide from San Simeon (35.6525N, 121.2417W), California. The individual was euthanized in MS-222 (1g/L), dissected to remove internal organs, decapitated, and preserved in liquid nitrogen. We used 1.21g of skin and muscle tissue to extract genomic DNA using the Genomic DNA and RNA purification kit provide by Macherey-Nagel (Düren, Germany) with three NucleoBond^®^ AXG columns. After extraction, the DNA samples were sheared with a 1.5 blunt end needles (Jensen Global, Santa Barbara, CA, USA) and run through pulse field gel electrophoresis, in order to separate large DNA molecules, for 16 hours. Samples were checked with a spectrophotometer (Synergy H1 Hybrid Reader, BioTek Instruments Inc., Winooski, VT) and Qubit^®^ fluorometer for quantification of our genomic DNA. We implemented two methods of high throughput sequencing techniques, Pacific Biosciences (PacBio) and Illumina platforms. For PacBio sequencing, genomic DNA was sized selected with BluePippin™ with a 15kb size cutoff, and 40 SMRT cells were sequenced with the PacBio RS II. In addition, from the same gDNA extraction, a multiplex gDNA-Seq Illumina sequencing library was prepared from size selected fragments which ranged from 500-700bp and sequenced on two lanes that resulted in short reads (100bp paired-end) on an Illumina HiSeq 2500. All genomic sequencing was completed at the University of California, Irvine (UCI) Genomics High-Throughput Facility (GHTF).

### Illumina and Pacbio Hybrid Assembly

We implemented multiple bioinformatic assembly programs to generate our final assembly of the *C. violaceus* genome with the following order of programs (Supplementary Figure S1; Supplementary Table S2). All computational and bioinformatics analyses were conducted on the High Performance Computing (HPC) Cluster located that University of California, Irvine. Sequence data generated from two lanes of Illumina HiSeq 2500 were concatenated and raw sequence reads were assembled through platanus v1.2.1 (Kajitani et al. 2014), which accounts for heterozygous diploid sequence data. Parameters used for platanus was -m 256 (memory) and -t 48 (threads) for this initial assembly. Afterwards, contigs assembled from platanus and reads from 40 SMRT cells of PacBio sequencing were assembled with a hybrid assembler dbg2olc v1.0 (Ye et al. 2014). We used the following parameters in dbg2olc: k 17 KmerCovTh 2 MinOverlap 20 AdaptiveTh 0.01 LD1 0 and RemoveChimera 1. Without Illumina sequence reads, we also conducted a PacBio reads only assembly with Falcon (https://github.com/PacificBiosciences/FALCON) with default parameters in order to assemble PacBio reads into contiguous sequences. The parameters we used were input type = raw, length_cutoff = 4000, length_cutoff_pr = 8000, with different cluster settings 32, 16, 32, 8, 64, and 32 cores, concurrency setting jobs were 32, and the remaining were default parameters. After our falcon assembly, we used the outputs from falcon and dbg2olc as input for quickmerge v1.0 (Chakraborty et al. 2016), a metassembler and assembly gap filler developed for long molecule-based assemblies. Several different parameters in quickmerge v1.0 were conducted as suggested by the authors until an optimal assembly was obtained with HCO 10 C 3.0 -l 2400000 -ml 5000. Afterwards, we polished our assembly with two rounds of QUIVER (Chin et al. 2013) and then finally using PILON v1.16 (Walker et al. 2014) to make improvements to the final genome assembly. N25, N50, and N75 was estimated with a perl script (Joseph Fass - http://bioinformatics.ucdavis.edu) from our final genome assembly of *C. violaceus*. With the final genome assembly, we processed our genome through repeatmasker v.4.0.6 (Smit et al. 2015) to mask repetitive elements with the parameter -species teleostei.

### Genome Size Estimation and Quantitative Measure of Genome Assembly

The c-value has been estimated for *Cebidichthys violaceus* (Hinegardner and Rosen, 1972), which is 0.81. Based on this c-value, the estimate of the genome size is ~792 Mb. In addition, we estimated the genome size using only Illumina sequences by using JELLYFISH v2.2.0 (Mar9ais and Kingsford, 2011). We selected multiple k-mers (25, 27, 29, 31) for counting and generating a histogram of the k-mer frequencies. We used a perl script (written by Joseph Ryan) to estimate genome size based on k-mer sizes and peak values determined from histograms generated in jellyfish.

We used tandem repeats finder (trf v4.07b; Benson, 1999) to identify tandem repeats throughout the unmasked genome. We used the following parameters in trf 1 1 2 80 5 200 2000 -d -h to identify repeats. Once the largest repeats were identified, we used period size of the repeats multiplied by the number of copies of the repeat to generate the largest fragments. This method was used to identify repetitive regions which can possibly represent centromere or telomere regions of the *C. violaceus* genome.

We used busco v3 (Simão et al. 2015) to estimate the completeness of our genome assembly with the vertebrata and Actinopterygii gene set [consists of 2,586 Benchmarking Universal Single-Copy Orthologs (BUSCOs)] to estimate completeness of our *C. violaceus* genome.

### RNA-Seq Tissue Extraction and Sequencing

Multiple individuals identified as *C. violaceus* were collected during the fall of 2015 for our transcriptomic analyses and annotation of the *C. violaceus* genome. The following IDs were selected for the five individuals for the transcriptomic analyses: CV96, CV97, CV98, CV99, and CV100. We selected nine tissues: brain, gill, gonads (testes), heart, liver, mid intestine, proximal intestine, pyloric caeca, and spleen. We extracted tissues from multiple *C. violaceus* individuals and preserved tissue in RNAlater^®^ (Ambion, Austin, TX, USA). Total RNA was extracted using a Trizol protocol, and an Agilent bioanalyzer 2100 (RNA nano chip; Agilent Technologies) was used to review quality of the samples. In order to produce a sufficient amount of total RNA from our extractions, we used a single or multiple *C. violaceus* individuals (up to four) to represent a specific tissue. We used the Illumina TruSeq Sample Preparation v2 (Illumina) kit along with AMPure XP beads (Beckman Coulter Inc.) and SuperScript™ III Reverse Transcriptase (Invitrogen) to prepare our tissue samples for Illumina sequencing. The following adaptor indexes were used for our analysis: brain - 6, gill - 2, gonadal tissue - 13, heart - 7, liver - 5, pyloic ceaca - 12, proximal intestine - 4, mid intestine - 14, and spleen - 15 (Supplementary Table S3). We size selected fragments that were on average 331 bp. In addition, we used a High Sensitivity Agilent Assay on an Agilent Bioanalyzer 2100, and Kappa qPCR for quantitative analyses before samples were sequenced. We multiplexed samples to 10 nM in 10 ul and were sequenced on two lanes on an Illumina HiSeq 2500 (100 bp Paired Ends) at UC Irvine’s GHTF.

### Transcript Assembly for all Tissues and Annotation

The following pipeline was used to assemble and measure expression of all transcripts from all nine tissue types (Supplementary Figure S2). Prior to assembly, all raw reads were trimmed with trimmomatic v0.35 (Bolger et al. 2014). Afterwards, trimmed reads were normalized using a perl script provided by trinity v r2013-02-16 (Grabherr et al. 2011). Prior to aligning transcriptomic reads to the genome, the final masked assembled genome was prepared with bowtie2-build v2.2.7 (Langmead and Salzberg, 2012) for a bowtie index and then all (normalized) reads from each tissue type were mapped using tophat v2.1.0 (Kim et al. 2013) to our assembled masked genome using the following parameters -I 1000 -i 20 -p 4. Afterwards, aligned reads from each tissue were indexed with samtools v1.3 (Li et al. 2009) as a BAM file. Once indexed through samtools, transcripts were assembled by using cufflinks v2.2.1 (Trapnell et al. 2012) with an overlap-radius 1. All assemblies were merged using cuffmerge and then differential expression was estimated with cuffdiff, both programs are part of the cufflinks package. All differential expression analyses and plots were produced in R (https://www.r-project.org/) using cummerbund tool located on the bioconductor website (https://www.bioconductor.org/). Once all transcripts were assembled, we ran repeatmasker v.4.0.6 with the parameter -species teleostei to mask repetitive elements within our transcriptomes.

All masked transcripts were annotated with the trinotate annotation pipeline (https://trinotate.github.io/), which uses Swiss-Prot (Boeckmann et al. 2003), Pfam (Finn et al. 2013), eggNOG (Powell et al. 2014), Gene Ontology (Ashburner et al. 2000), SignalP (Petersen et al. 2011), and Rnammer (Lagesen et al. 2007). We also processed our transcripts through blastx against the UniProt database (downloaded on September 26th, 2017) with the following parameters: num_threads 8, evalue 1e-20, and max_target_seqs 1. The blastx output was processed through trinity analyze_blastPlus_topHit_coverage.pl script to count the amount of transcripts of full length or near full length. To provide a robust number of full-length transcripts, the assembled genome was processed through AUGUSTUS v3.2.1 (Stanke et al. 2006) without hints using default parameters for gene predictions using a generalized hidden Markov model in order to identify genes throughout the genome, and predicted transcripts were also masked for repetitive elements through repeatmasker.

### Heatmap of all Tissues and Genes Associated with Diet Specialization

Differentially expressed genes for all tissue types were viewed with a heatmap that was generated with the cummerbund library (Supplementary Figure S3). Candidate genes which pertained to glycolytic, lipid metabolism/gluconeogenesis, ketone degradation, glucosidases (both α and β), proteases, and lipases were identified in the *C. violaceus* transcriptome by scanning the annotation of cufflinks assembled transcripts and used to generate our heatmap.

### Identification of Candidate Genes and Copy Number

Amylase and Bile Salt Activated Lipase are candidate genes of interest due to their properties of breaking down starch (α-glucans) and dietary lipids, respectively, and we were interested in identifying copy number of these two candidate genes. Previously published variants of *amy2* (*amy2a* and *amy2b*; German et al. 2016) found in *C. violaceus* which were deposited on NCBI (KT920438 and KT920439) were used as queries and searched throughout our assembled genome using both mummer v3.23 (Kurtz et al. 2004) and blast (Altschul et al. 1997) to identify gene copies. Afterwards, we used a perl script (Lawrence et al. 2015) to trim the contig which contained the amylase gene copies and neighboring loci and were viewed with the online version of AUGUSTUS v3.2.3 (Stanke and Morgenstern, 2005). Next, we used amylase sequences from multiple Stichaeid intertidal species that represent a broad spectrum of diet specializations (German et al. 2016; Fig. 1). We selected *Anoplarchuspurpurescens* (carnivore), *Dictyosoma burgeri* (carnivore), *Phytichthys chirus* (omnivore), *Xiphister atropurpureus* (omnivore), and *X. mucosus* (herbivore; German et al. 2016). Our predicted amylase sequences from *C. violaceus* and orthologous sequences from the five other intertidal prickleback species were aligned in mega v7.0.26 (Kumar et al. 2016) with muscle (with codons) and then selection was estimated using adaptive Branch Site REL (aBSREL) and Mixed Effects Model of Evolution (MEME) as well as signatures of recombination Genetic Algorithm for Recombination Detection (GARD) as part of the datamonkey 2.0 web application (Weaver et al. 2018).

To identify Bile Salt Activated (*BSA*) lipase in the *C. violaceus* genome, we used the haddock (*Melanogrammus aeglefinus;* AY386248.1) BSA lipase and BLASTed this coding gene against our assembled transcriptomes where the highest bit score and percent identity (greater than 70%) were used to identify the orthologs. Once BSA lipase transcripts were identified, then we used mummer and blast to identify gene copies of BSA lipase within the *C. violaceus* genome. Again, we used a perl script (Lawrence et al. 2015) to trim the contig which only contained the BSA lipase genes and neighboring loci and were viewed with AUGUSTUS. We identified orthologous sequences from assembled transcriptomes from *Xiphister mucosus*, *Xiphister atropurpureus*, *Anoplarchus purpurescens*, and *Phytichthys chirus* pyloric cacea samples, which were generated for another study (Herrera et al. unpublished). All sequences were aligned in mega v7.0.26 and then selection and recombination were estimated in datamonkey server v2.0 with aBSREL, MEME, and GARD.

### Identification of orthologs across teleost fishes

All *C. violaceus* transcripts predicted from AUGUSTUS were used for identifying orthologs among ensembl protein datasets of bony (teleost) fishes and a lobed finned fish (coelacanths). Then the following teleost and non-teleost genomes were taken from ensembl release 89 for our comparative analysis of orthologs and phylogeny of fishes; *Poecilia formosa* (Amazon molly), *Astyanax mexicanus* (blind cave fish), *Gadus morhua* (Atlantic Cod), *Takifugu rubripes* (Fugu), *Oryzias latipes* (Japanese medaka), *Xiphophorus maculatus* (platyfish), *Lepisosteus oculatus* (spotted gar), *Gasterosteus aculeatus* (stickleback), *Tetraodon nigroviridis* (green spotted puffer), *Oreochromis niloticus* (Nile tilapia), *Danio rerio* (zebrafish), *Latimeria chalumnae* (coelacanth). All *C. violaceus* transcripts were translated into protein sequences by using ORFPREDICTOR (Min et al. 2005). From all 13 fish species, only sequences with 60 amino acids or longer were used for our analyses. We conducted a pairwise identification of orthologs by using INPARANOID v4.0, (O’brien et al. 2005) in which we conducted 78 possible pairwise comparisons, where N is number of taxa [(N(N-2))/2= possible pairwise comparisons]. From the outputs of inparanoid, we used quickparanoid (http://pl.postech.ac.kr/QuickParanoid/) to identify orthologous clusters from all 13 species. Once single copy orthologs were identified from all datasets, we used custom python scripts to align sequences using MUSCLE (Edgar, 2004) and then a consensus phylogenetic tree was generated using PhyML v3.1 (Guindon et al. 2010) with 1,000 bootstrap replicates based on the lowest average gaps present in our ortholog cluster alignments of concatenated 30 protein coding genes from all 13 taxa.

### Comparative Analysis for Syntenic Regions across Teleosts fishes

We selected the following fish genomes: zebrafish (*Danio rerio*), stickleback (*Gasterosteus aculeatus*), spotted gar (*Lepisosteus oculatus*), and the Japanese medaka (*Oryzias latipes*) to identify syntenic regions with our *C. violaceus* genome assembly. All genomes were masked using repeatmasker v.4.0.6 with the parameter -species teleostei. To identify syntenic regions we used contigs from our *C. violaceus* which represent 1MB or larger and then concatenated the remaining contigs. We used satsuma v3.1.0 (Grabherr et al. 2010) with the following parameters -n 4 -m 8 for identifying syntenic regions between zebrafish, stickleback, spotted gar, Japanese medaka and the *C. violaceus* genome. Afterwards, we developed circos plots to view syntenic regions shared between species using CIRCOS v0.63-4 (Krzywinski, 2009).

## Supplementary results and discussion

### I - Genome and Transcriptome Assembly

From PacBio sequencing, we were able to generate ~29,700 Mb of sequence data from 40 SMRT cells, this represents ~37X coverage based on the c-value estimated for *C. violaceus* (792 Mb; Hinegardner and Rosen, 1972). From two lanes of Illumina sequencing we were able to generate 84,539 Mb which represents ~107X coverage of the genome. From the platanus assembly with Illumina only sequence data, our N50 was 2,760 basepairs. When combining platanus assembled contigs and 40 SMRT cells of PacBio for a hybrid assembly in dbg2olc, we managed to obtain an N50 of 2.21Mb. Afterwards, when using FALCON (PacBio only reads) we managed to obtain an N50 of 2.45Mb. We used the falcon assembly and the dbg2olc assembly and through quickmerge, we obtained an N50 of 6.69 Mb. Following two rounds of quiver, and pilon, we assembled our final draft genome which composed of 467 contigs, and obtained N25, N50, and N75 values of 15.17, 6.71, and 1.85 (Mb) respectively which were composed of 593,001,491 base pairs (Supplementary Figure S4). Genomic regions that were masked in repeatmasker were masked and the identities of the repetitive elements are present in Supplementary Table S4.

### II - Estimation of Genome Size, Completeness Assessment, and Tandem Repeats Throughout the Genome

By using jellyfish we estimated the genome size based on an average of four k-mer size counts (25, 27, 29, 31) 656,598,967 base pairs based with a standard deviation of 4,138,853 base pairs (Supplementary Figure S5; Supplementary Table S5). Through BUSCO v3 (, there were 97% (2,508 genes out of 2,586) complete orthologs detected which included 1.3% duplicated orthologs. In addition, there were 1.1% (28 BUSCOs) partial orthologs present and 1.9% (50 BUSCOs) of orthologs were not detected in the *C. violaceus* genome (Supplementary Table S6).

In identifying large genomic regions of tandem repeats we were able to identify about 38,448 repeated loci where the repetitive sequence (period) range from 1 to 1,983 and amount of repeats identified were 1.8 to 14,140.8bps. The largest repeat locus (period size multiplied by the repeat amount) was 109,161bps with a period of 90bp and repeat amount of 1,212.9bps. We saw an increase in size for 35 loci which had repeat locus the size of 32,594.1bps and greater (Supplementary Figure S6-7) as compared to any other repetitive locus.

### III – Assembly and Annotation of Nine Tissue Transcriptomes

From five individuals that we selected for our transcriptomic analyses, the total reads mapped back to the genome ranged from 67.37% (liver) to 84.8% (heart; Supplementary Table S2). The range of transcripts present in each tissue type ranged from 20,008 (liver) to 78,629 (gill) transcripts.

From the tuxedo package, there were 101,922 transcripts estimated from the nine tissues. When evaluating Fragments Per Kilobase per Million mapped Reads (FPKM) for each of the nine transcriptomes, we see the lowest median with the liver and the highest median with the gill tissue (Supplementary Figure S8). All other tissues appeared to have a similar profile. In addition, we see that the gill tissue had a short Quartile group 2 as compared to the other tissue types (except liver tissue; Supplementary Figure S8). When we look at the differentially expressed genes across all tissue types, we see that there are cluster of genes highly expressed in the liver as compared to some of the tissues (Supplementary Figure S9). By using getSig in the cummerbund package we evaluated which genes are significantly regulated, with the highest number (1,383) between liver and brain, as these tissues are highly specialized. When we look at Jensen-Shannon Distances (Supplementary Figure S10-11), we see that pyloric caeca and middle intestine have similar expression profiles, brain and gonad have similar profiles, and the proximal intestine has a very different expression profile.

From the assembled transcripts in trinity and predicted transcripts from AUGUSTUS, there were a total of 105,167,222 and 44,120,550 bases with 0.08% and 2.15% of bases masked respectively (Supplementary Tables S7-8). When annotated, there were 65,535 transcripts in trinotate and there were only 26,356 transcripts with a blastx hit identified. When conducting a blastx on our transcripts to identify full length transcripts in our dataset, we were only able to obtain 5,199 transcripts which had a 80% hit coverage (Supplementary Tables S9-10). When using only using augustus (without hints) and identified 29,525 genes. There were 29,485 genes which had 60 amino acids or greater in our transcriptomic dataset (Supplementary Table S8).

### IV - Candidate Genes for Digestion and Metabolism

We were able identify candidate genes associated with digestion, fermentation, ketone degradation in our transcriptomic assembly (Fig. 3A & B) and view differential gene expression patterns across the nine tissues where we have transcriptomic dataset (Supplementary Figure S3). We were not able to distinguish between *amy2a* and *amy2b* in our transcriptomic assembly. Therefore, we used the augustus gene prediction from the genome as a transcriptome reference to detect the *amy2a* and *amy2b* genes and mapped our transcriptome reads back to this reference transcriptome dataset. From this dataset, we were able to detect *amy2a* and *amy2b* gene expression profiles. As expected, we see high expression profiles for genes associated with digestion and metabolism in the pyloric ceca, proximal intestine, mid intestine, and liver (Fig. 3A & B).

### V - Amylase and Bile Salt Activated Lipase

From our mummer and blast search for pancreatic amylase we have identified three tandem copies of amylase (*amy2a*) and (*amy2b*), as opposed to the six haploid copies detected in the German et al. 2016. We identified amylase on contig 440 and we see two hypothetical proteins between the three amylase genes and a transposase near *amy2b* (Fig. 4B; Supplementary Figure 12). In addition, each amylase gene is preceded by a 4.3K20bp DNA element encoding a transposase (Fig. 4B, Supplementary Figure 13). The three tandem amylase loci differs from the estimated six haploid copies (based on gene dosage curves using RT-qPCR) proposed to be present in the *C. violaceus* genome by German et al. (2016). Upon further inspection, we have reached the conclusion that the per cell gene count of the German et al. (2016) study is the diploid copy number, not the haploid copy number (C. *violaceus* is a diploid, vertebrate). Hence, the copy number based on gene dosage curves is three for amylase in general, with roughly two copies for *amy2A* and one copy for *amy2B* (German et al. 2016) which agrees completely with what is observed in our genomic assembly. Although there is the possibility for copy number variation amongst individuals within a population, as there is for human salivary amylase (Perry et al. 2007) and dog pancreatic amylase (Axelsson et al. 2013), that is not what was observed by German et al. (2016), as that would entail different methodology and more robust sampling of *C. violaceus* individuals.

We only selected one copy of *amy2a* because they are identical and *amy2b* when estimating selection in datamonkey. When testing all 11 branches for seven taxa in aBSREL, we see only one branch under episodic diversifying selection (C. *violaceus, amy2b*) with a significant uncorrected p-value of 0.0044 (Fig. 4C). We do not see this pattern of positive selection in any of the other branch with a significant p-value. In MEME, we that three episodic positive selection with a p-value under 0.05 (sites: 41, 256, and 279; Fig. 4D) and in GARD there was no evidence of recombination.

We focused on the loci that encode for Bile Salt Activated Lipase and were able to identify the four tandem copies of BSAL on contig 445 (Fig. 5B; Supplementary Figure S14).

We estimated selection in absrel with 11 taxa and found three branches out of 19 to be under episodic diversifying selection (Fig. 5C). These three branches, *Cviolaceus_2* (uncorrected p-value 0.0008), Node 9 (uncorrected p-value 0.0000), and *Xmucosus_2* (uncorrected p-value 0.0000). In addition, we see 14 sites under episodic positive selection with the following sites under selection and a p-value less than 0.05: site 92, 94, 115, 139, 144, 151, 156, 160, 356, 429, 436, 443, 460, and 475 (Fig. 5D). In addition, we see evidence of recombination in BSAL and with a p-value of 0.01 we see 1 breakpoint with significant topological incongruence at site 534 when we used GARD. We also used the Bile salt-activated lipase (carboxy ester lipase) from *Homo sapiens* to align with our *C. violaceus* sequences and the other four intertidal prickleback species (Supplementary Figure S15).

With regards to elevated lipase activity in the taxa consuming more fiber in their diets, it is known that in industry settings, adding microcrystalline fiber to lipolytic reactions acts to stabilize the lipase proteins and can increase lipase activities (Cai et al. 2018; Kim et al. 2017) However, these fibrous compounds added directly to the reactive environment. In our case, when measuring lipase activities, we homogenize the tissues separate from gut contents, and then centrifuge the homogenates to get the supernatant, which would contain only those enzymes that are soluble and not bound to larger molecules, like fiber. Hence, the elevated lipase activities measured *in vitro* in our investigations cannot be coming from the potential effects of fiber on the lipase proteins themselves, even if these interactions might act to aid lipolytic action *in vivo* within the gut environment. We are, therefore, confident that the increased lipase activities in the algae-eating fishes are due to the molecular differences in these proteins (Fig. 5), and any differences in gene expression.

### VI - Orthologs and Phylogenetic Analyses

We constructed a phylogenetic tree using maximum likelihood using thirteen fish taxa which including the *Cebidichthys violaceus* which included 30 loci and 33,508 bases with 1,000 bootstrap replicates (Fig. 1). We used jmodeltest v2.1.0 and with AICc we detected that GTR+I+G was the best model selected for our phylogenetic analyses. All 30 loci used for our phylogenetic analyses were extracted from orthologs detected in inparanoid (Supplementary Table S11).

### VII - Syntenic regions across multiple fish species

We have 114 contigs which have a 1MB or greater, and we pooled all contigs which had less than 1MB (353 contigs) were merged together in our synteny analyses. When we compare our assembled genome to the *G. aculeatus* genome, we see multiple homologous regions between the two species, and multiple loci from each linkage group of the *G. aculeatus* genome represented in the *C. violaceus* genome; Supplementary Figure S16). When comparing the *L. oculatus, D. rerio*, and *O. latipes* genomes to our *C. violaceus* genome (Supplementary Figure S17-S19), we also see each chromosome/linkage group represented in the *C. violaceus* genome and strong synteny between *O. latipes* and *C. violaceus* whereas we less homologous strands between the *C. violaceus* genome and the *D. rerio* or *L. oculatus* genomes.

### VIII - Opsin Gene Copies and Selection

After reviewing our busco analyses for gene duplicates present in our *C. violaceus* genome, we identified three Opsin Short Wave Sensitive (*Opn1sw*) genes in tandem on contig 443 (Supplementary Figure S20). We find this interesting because *C. violaceus* endures a period of time out of water during low tide, in which Horn and Riegle (1981) showed that a large *C. violaceus* (~24 cm SL; 92 g) can survive out of water for 37 hours. The ability to survive out of water may require adaptations of vision when exposed to air during low tides, in which we further evaluated the *Opn1sw* gene copies for signatures of positive selection. In addition, we identified and compared gene copy numbers of short wave opsin genes from *D. rerio, Oreochromis niloticus*, and *G. aculeatus* genomes, which have one or two gene copies present (Supplementary Figure S20). In addition, we estimated selection by using the datamonkey server v2.0 by using absrel and MEME. We observed one branch of episodic diversifying selection out of 11 leading to opn1sw2a along with an opn1sw2 identified in *G. aculeatus* with an uncorrected p-value of 0.0028. In addition, we detected episodic positive/diversifying selection at 6 sites. The sites under selection and a p-value less than 0.05 were the following: site 4, 95, 165, 202, 224, and 337 (Supplementary Figure S20). From this analysis, evaluation of *Opn1sw* gene sequences from subtidal and intertidal Stichaeids can elucidate how vision may play an important role for intertidal prickleback species.

**Supplementary Figure S1:**
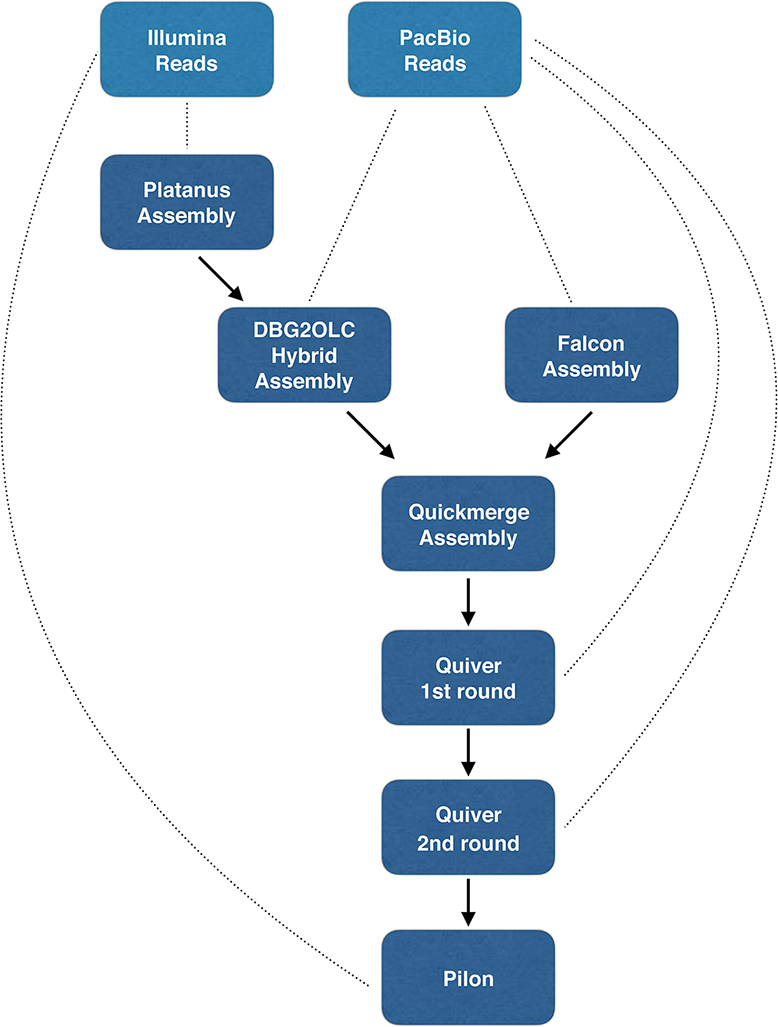
Flowchart of our final genome assembly from Illumina (two lanes of PE 100bp) and Pacific Biosciences (40 single molecule real-time [SMRT] cells) sequence reads. Light blue boxes indicate raw sequence reads and dark blue boxes indicate bioinformatic programs used for the assembly. (…………) indicates the type of sequence information that was used for the assembly method. Arrows indicate the next step taken to proceed in the genome assembly of *Cebidichthys violaceus*.

**Supplementary Figure S2:**
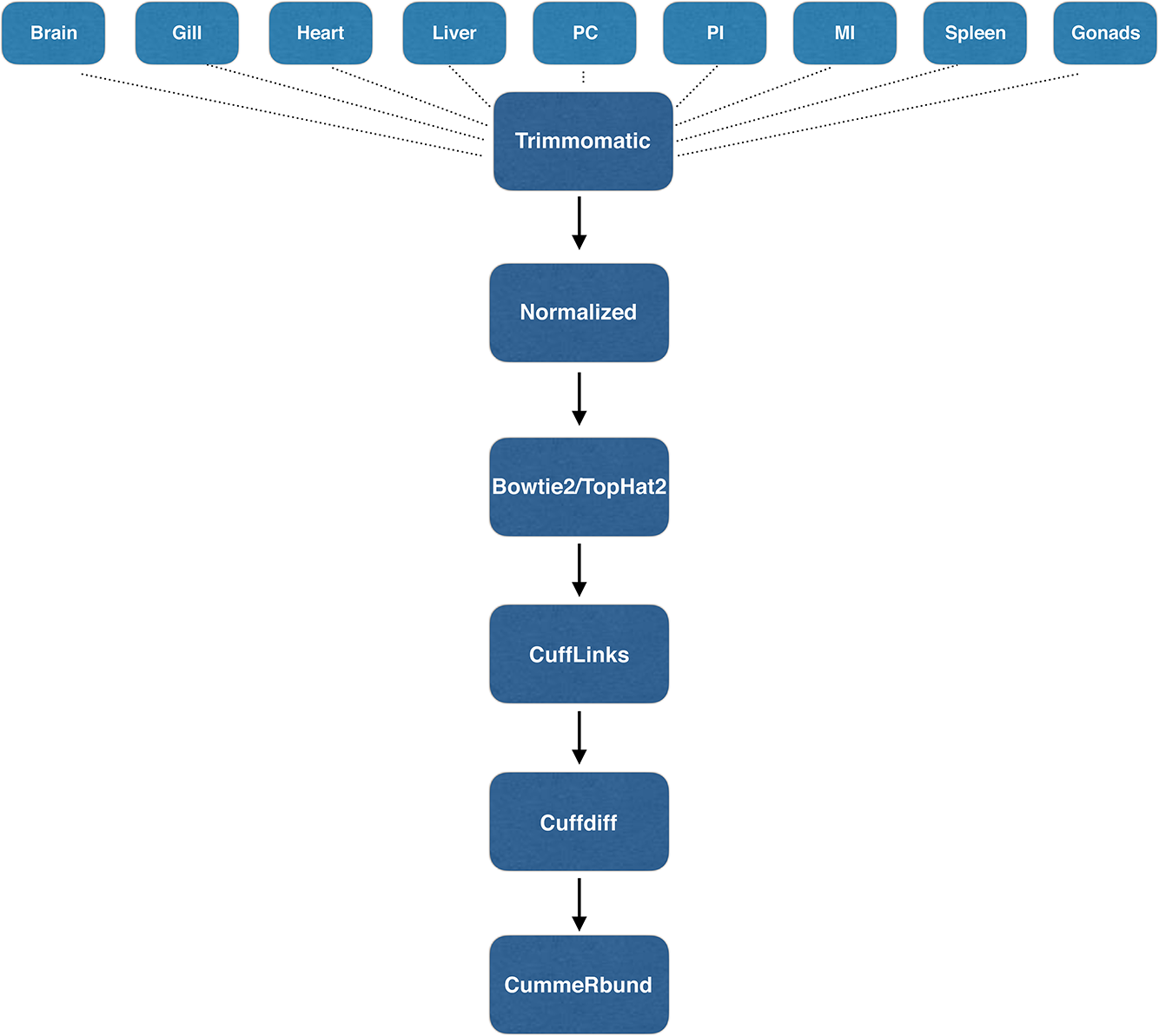
Flowchart of our genome guided transcriptome assembly from nine tissues using Illumina (two lanes of PE 100bp). Light blue boxes indicate raw sequence reads for nine tissues: liver, heart, gill, pyloric caeca (PC), proximal intestine (PI), middle intestine (MI), spleen, gonad (testes), and brain. Dark blue boxes indicate the bioinformatic program used in the pipeline for for trimming/cleaning reads, normalizing reads, assembling transcripts with our assemble genome as a reference, and estimating differential gene expression (DEGs) and analysis of DEGs. (…………) indicates the type of sequence information that was used for the start of the assembly method. Arrows indicate the next step taken to proceed in the transcriptome assembly and analysis.

**Supplementary Figure S3:**
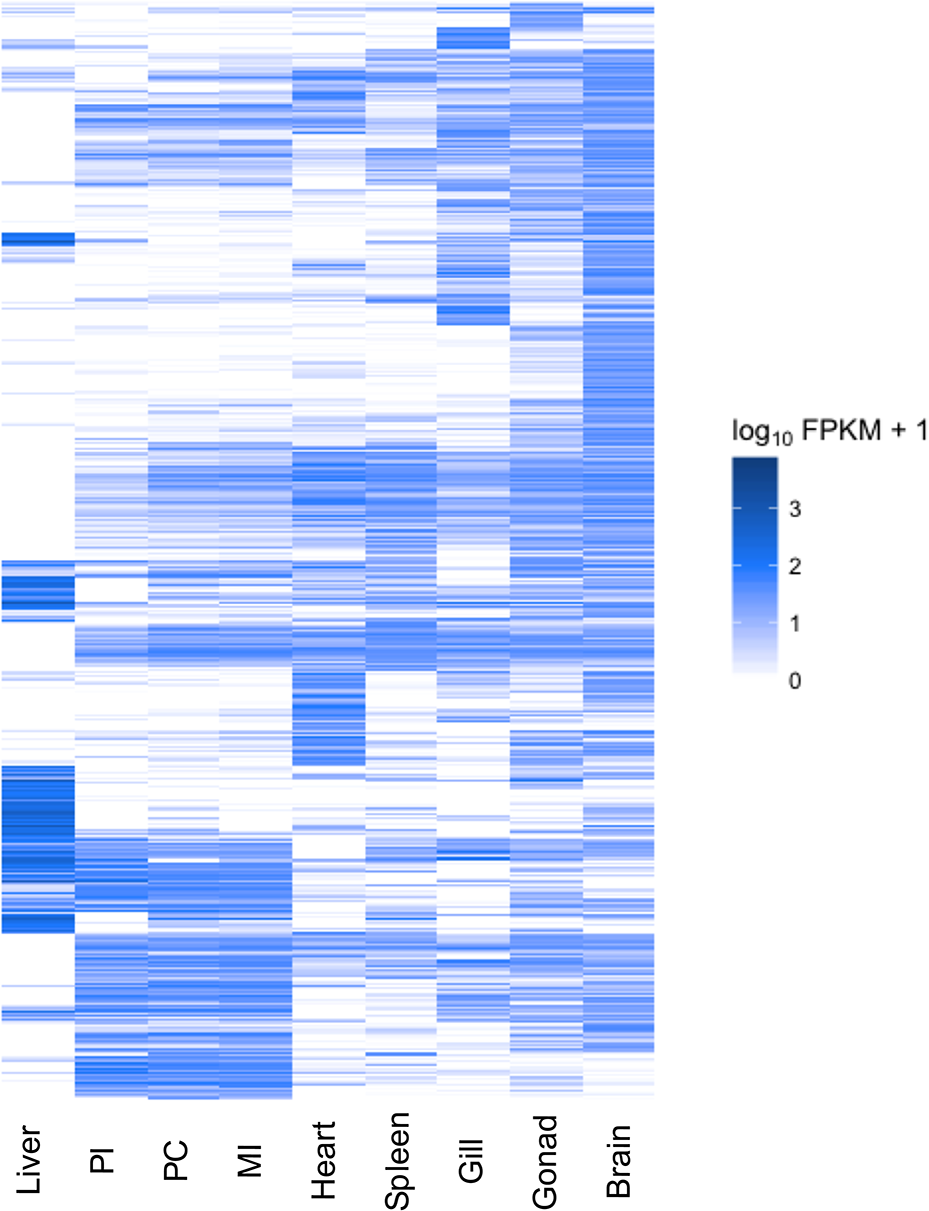
Heatmap showing Differentially Expressed Genes for nine tissues of *Cebidichthys violaceus*. Heatmap was generated with the csHeatmap feature in cummerbund, where dark blue represents a high FPKM value and white indicates a low FPKM value. There were 15,490 differentially expressed genes across all nine tissue types.

**Supplementary Figure S4:**
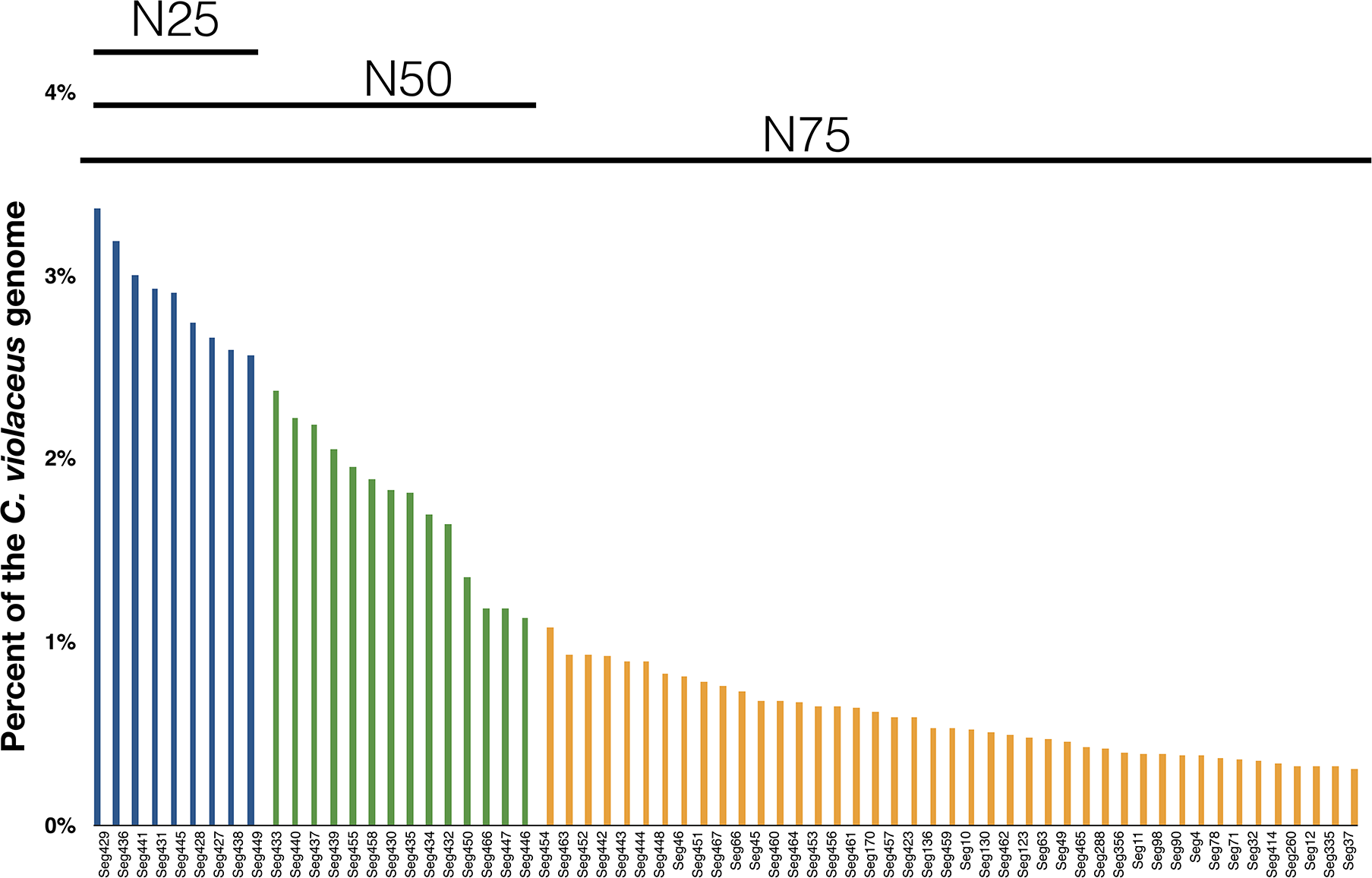
Estimation of completeness of the *Cebidichthys violaceus* genome. N25 (blue bars), N50 (blue + green bars), and N75 (blue + green + orange bars) values was estimated for the C. violaceus genome. There are 66 contigs which represent the N75 of the C. violaceus genome. Contig ID is labeled along the x-axis.

**Supplementary Figure S5:**
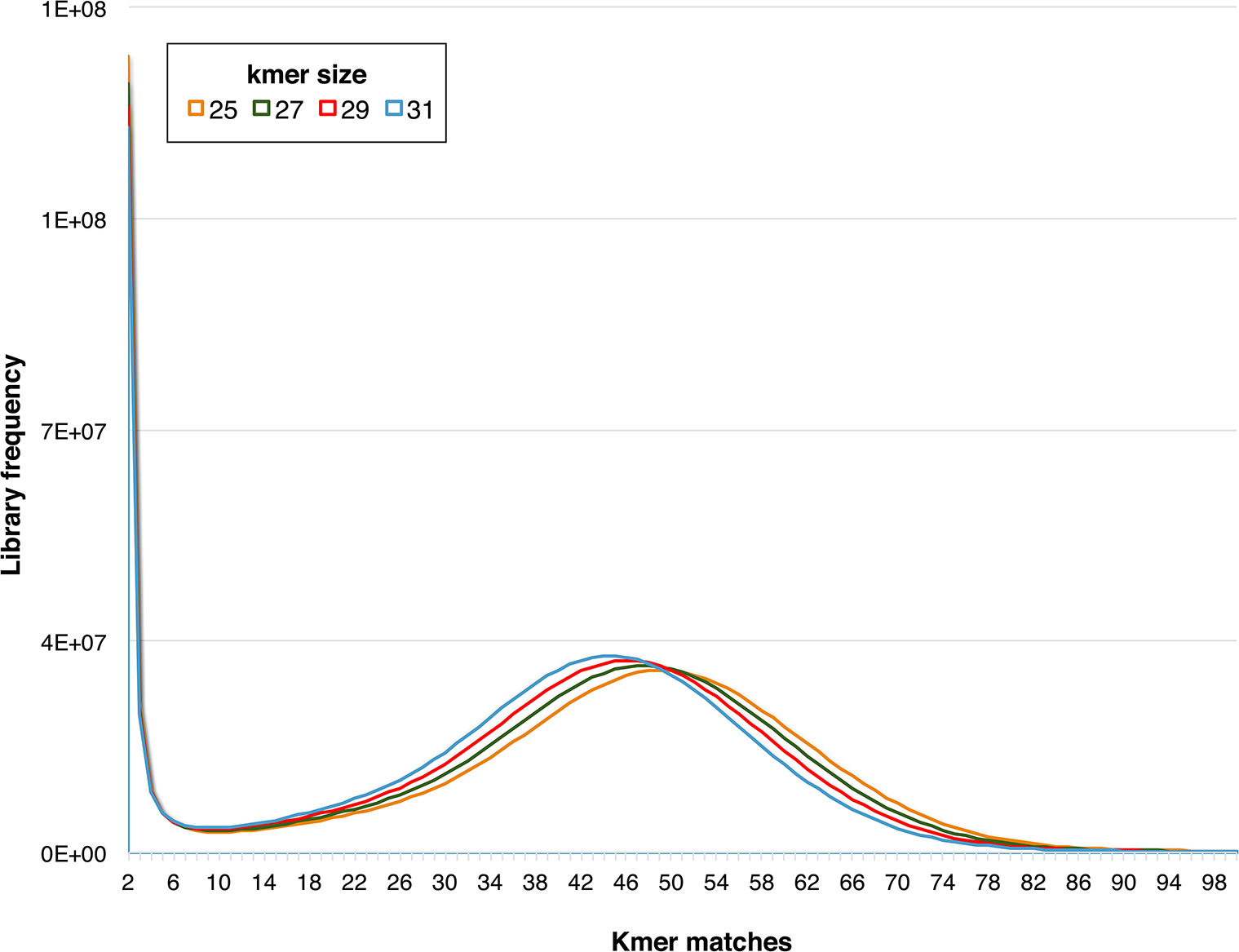
K-mer frequency for *Cebidichthys violaceus*. Use of raw Illumina (only) reads from *C. violaceus* gDNA to estimate the *C. violaceus* genome size. K-mer sizes of 25, 27, 29, and 31 were selected to generate histograms.

**Supplementary Figure S6:**
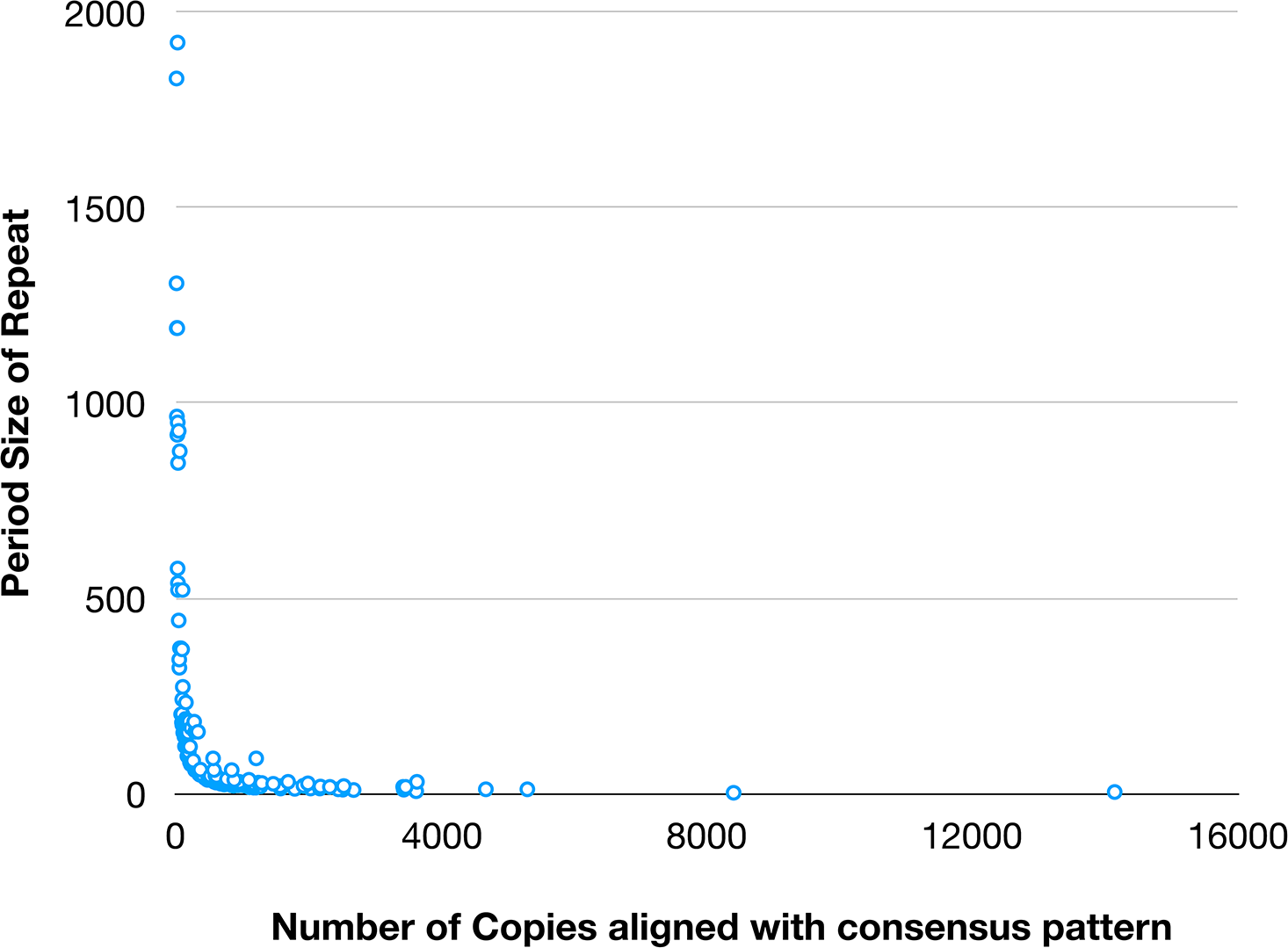
Period size and copies of pattern of tandem repeats identified in the *Cebidichthys violaceus* genome. The length of the tandem repeat sequence and the size of the repeat found.

**Supplementary Figure S7:**
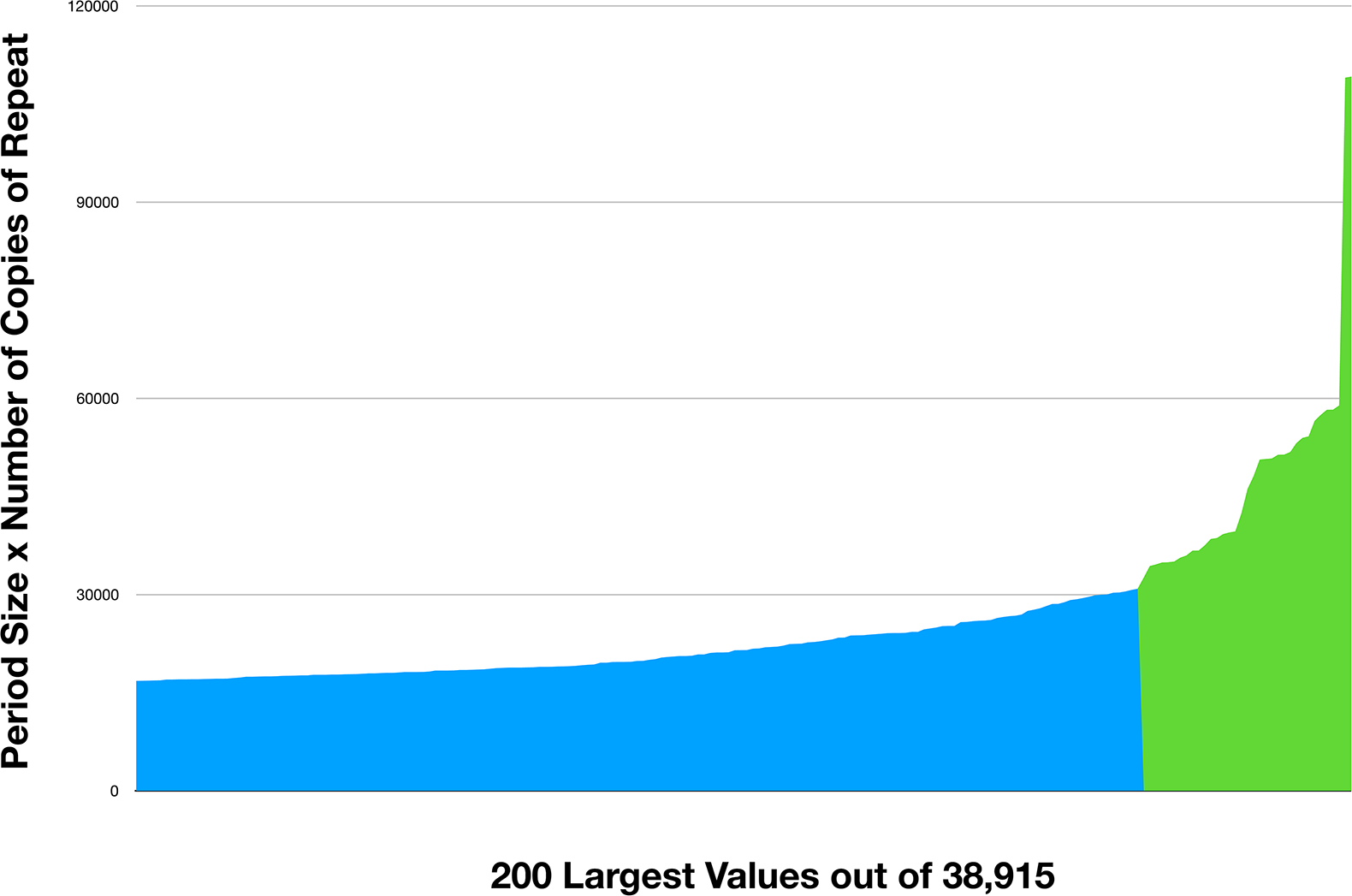
Histogram of period size multiplied by number of copies of repeat. Total of the 200 longest repeats identified, where blue indicates values of 30,877 base pairs (bps) or less. Green indicates values greater than 30,877 bps.

**Supplementary Figure S8:**
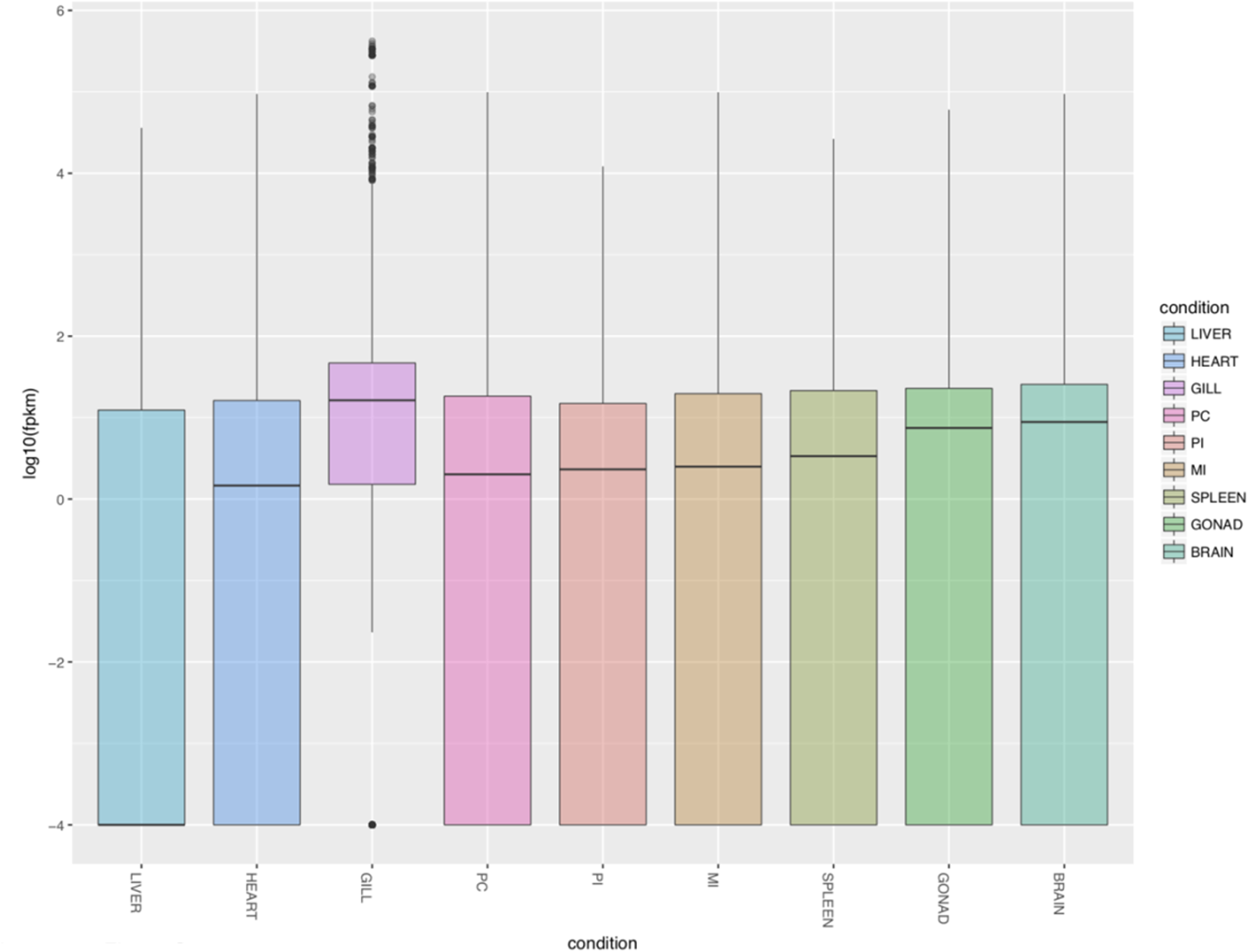
Boxplots visualized for all nine tissue types displaying summary statistics of Fragments Per Kilobase of transcript per Million mapped reads (FPKM). Boxplots were generated with the csBoxplot function in cummerbund for nine tissues.

**Supplementary Figure S9:**
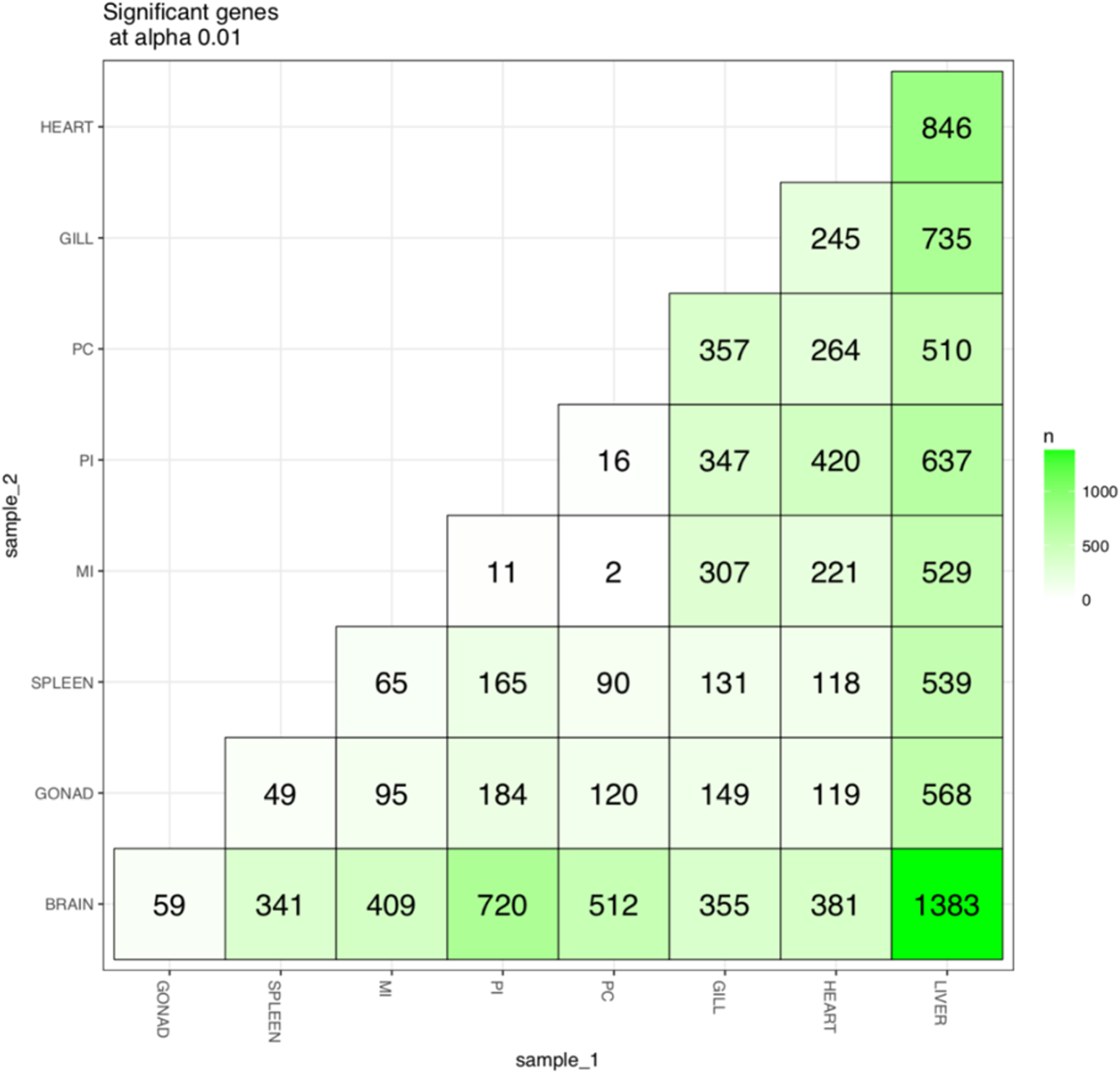
Significant features (pairwise) for all nine tissue types. Significant features were estimated by using the sigmatrix function in CUMMERBUND. The darker green shades indicate a higher significant feature identified, whereas a lighter green/white shade indicates a less significant feature identified.

**Supplementary Figure S10:**
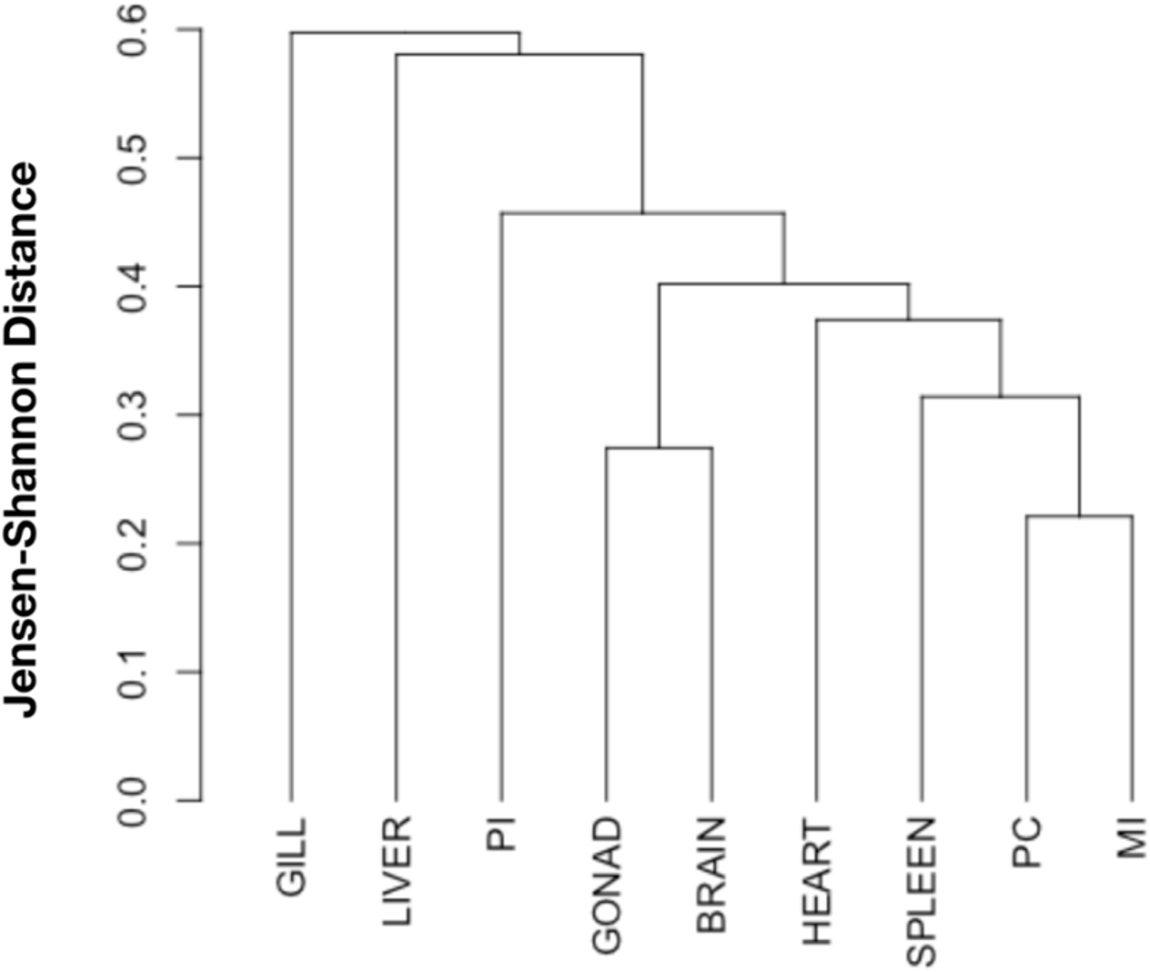
Dendrogram of all nine tissue transcriptomes to determine relationships of each tissue type. A dendrogram was constructed of all nine tissue types by using Jensen-Shannon (JS) distances as shown.

**Supplementary Figure S11:**
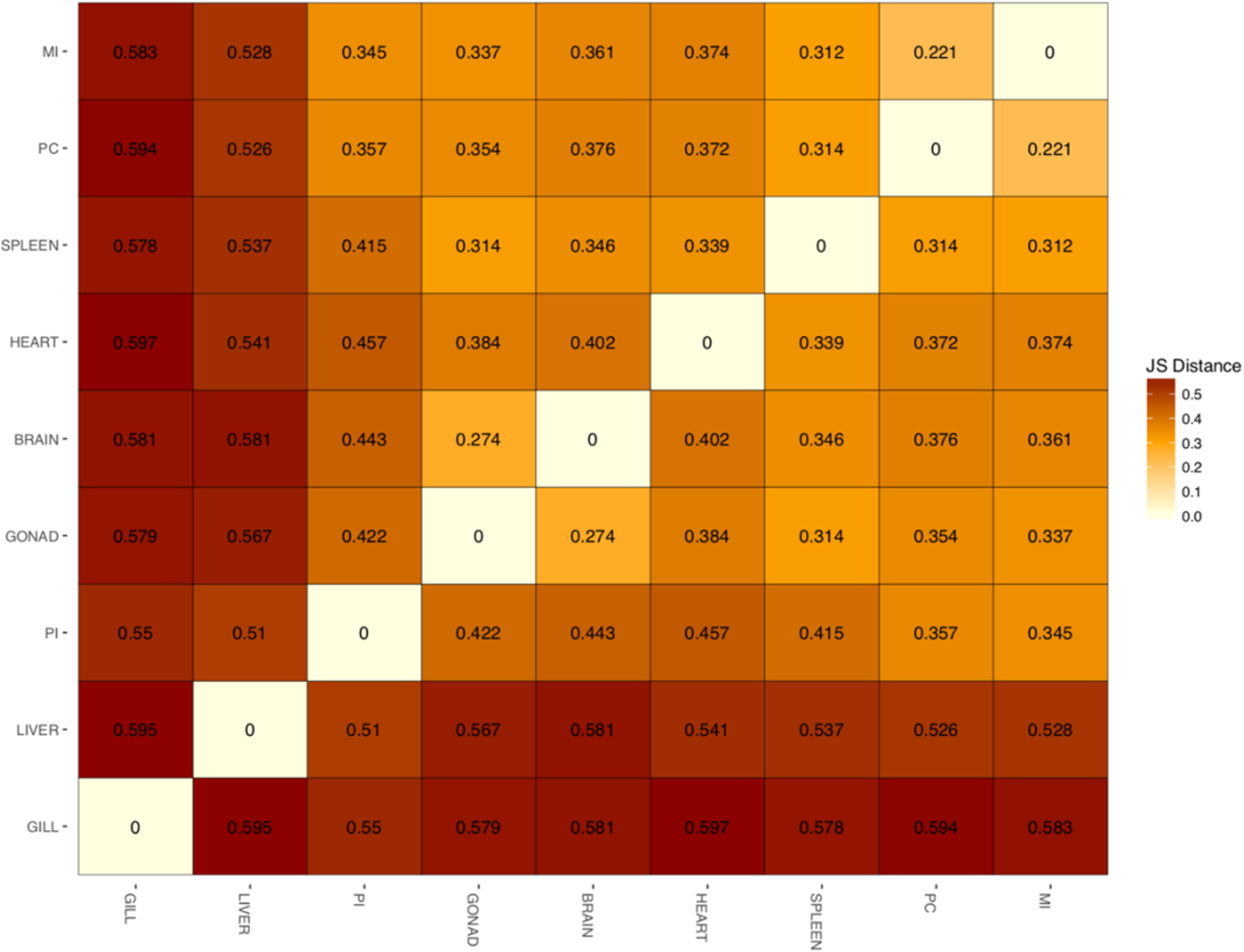
Distance matrix of all nine tissue transcriptomes based on Jensen-Shannon (JS). Plot was constructed with csDistHeat function in cummerbund. Dark red indicates an increased JS distance between the pairwise comparison. Lighter red/white indicates less distance between the pairwise comparison.

**Supplementary Figure S12:**
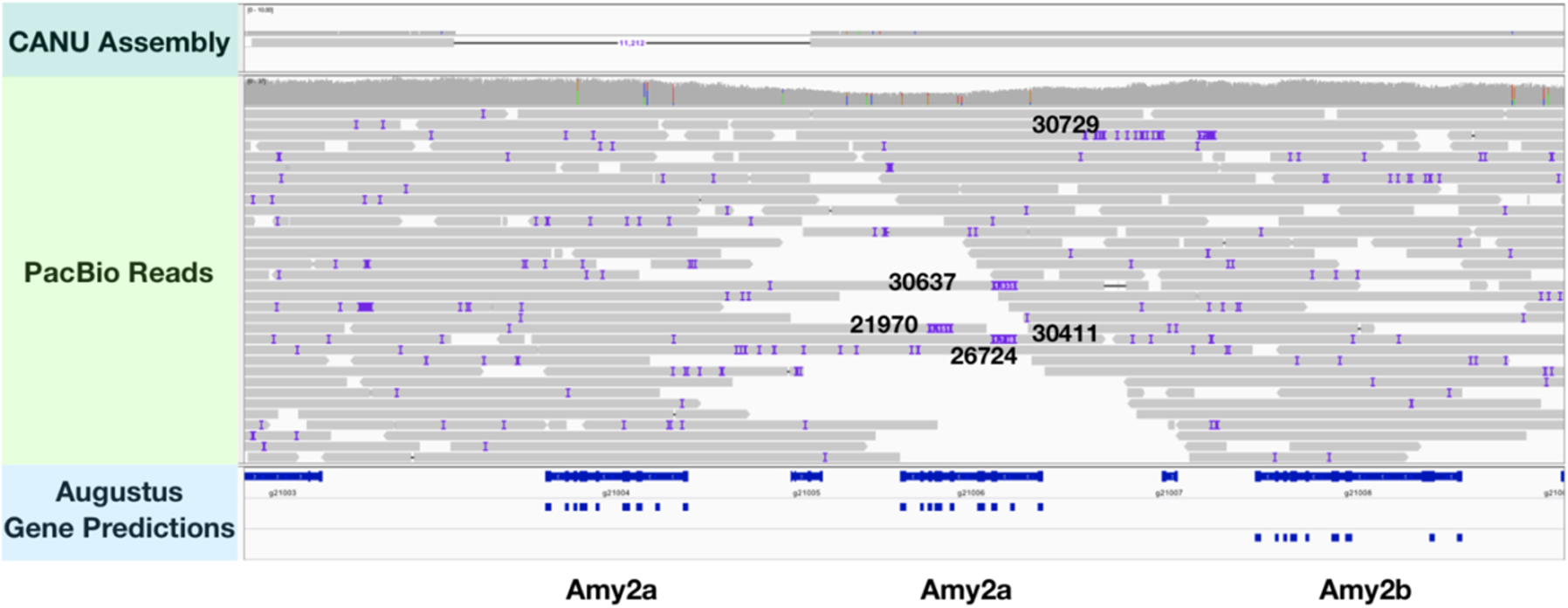
Pacific Biosciences (PacBio) reads mapped to C. violaceus genome assembly. PacBio reads with read ID numbers labeled which span regions of the three amylase loci (two amy2a loci and the amy2b) on contig 440. augustus gene predictions are used to reference where amy2 loci are located on the C. violaceus genome.

**Supplementary Figure S13:**
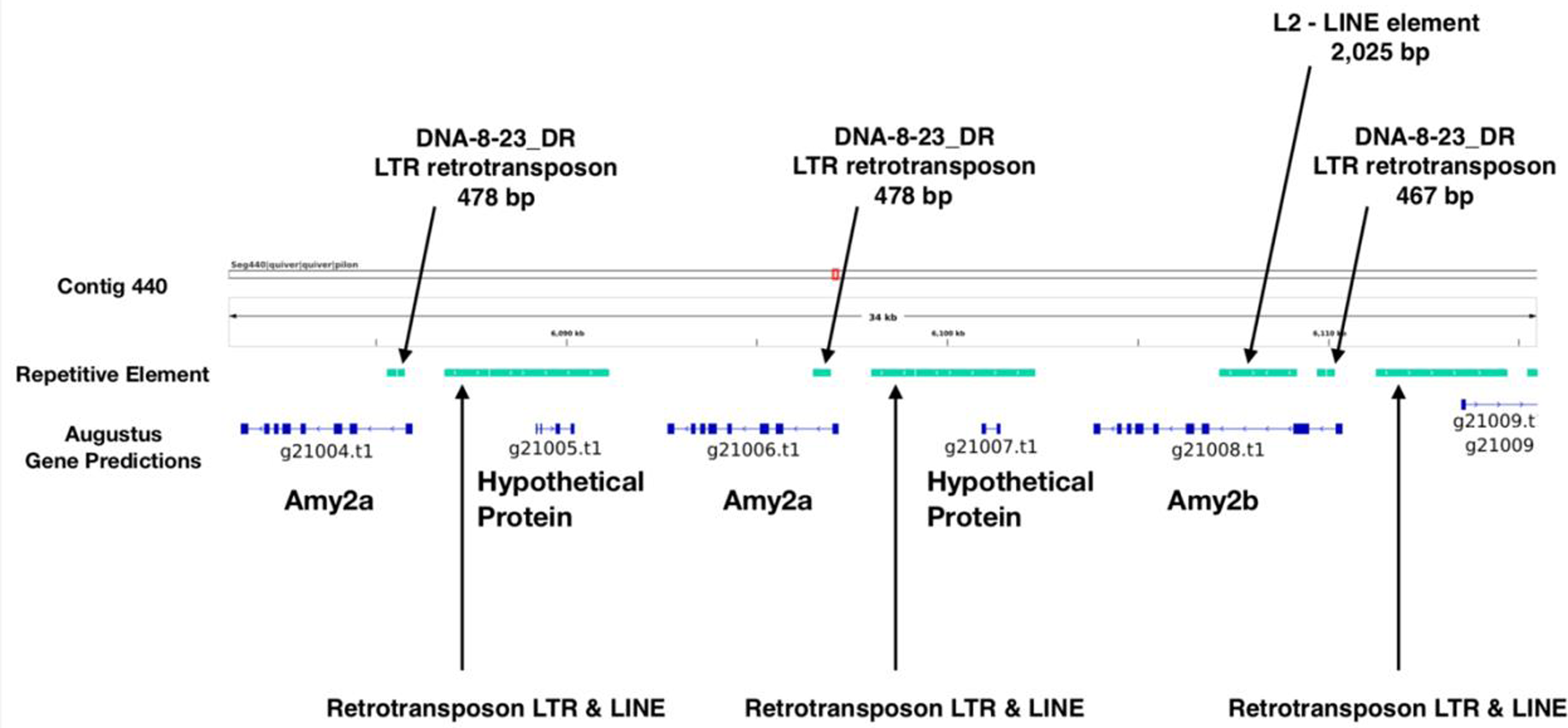
Repetitive elements identified adjacent to Amylase loci. Light green bars are repetitive elements identified on contig 440 and blue bars represent exons of amylase loci (augustus gene prediction).

**Supplementary Figure S14:**
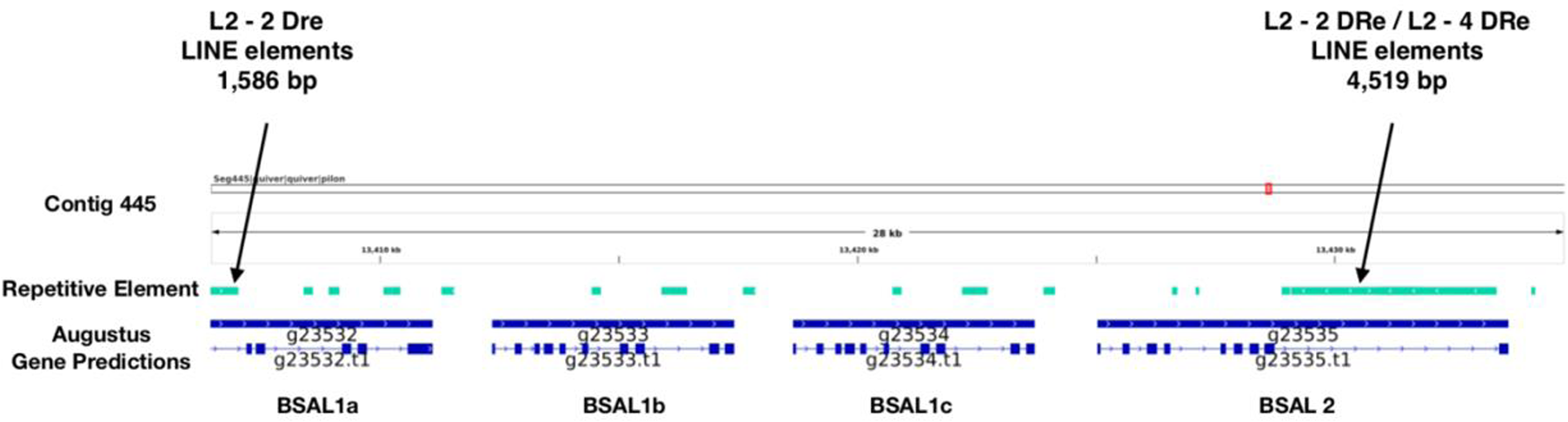
Repetitive elements identified adjacent to Bile Salt-Activated Lipase (BSAL) loci. Light green bars are repetitive elements identified on contig 445 and blue bars represent exons of BSAL loci (augustus gene prediction).

**Supplementary Figure S15:**
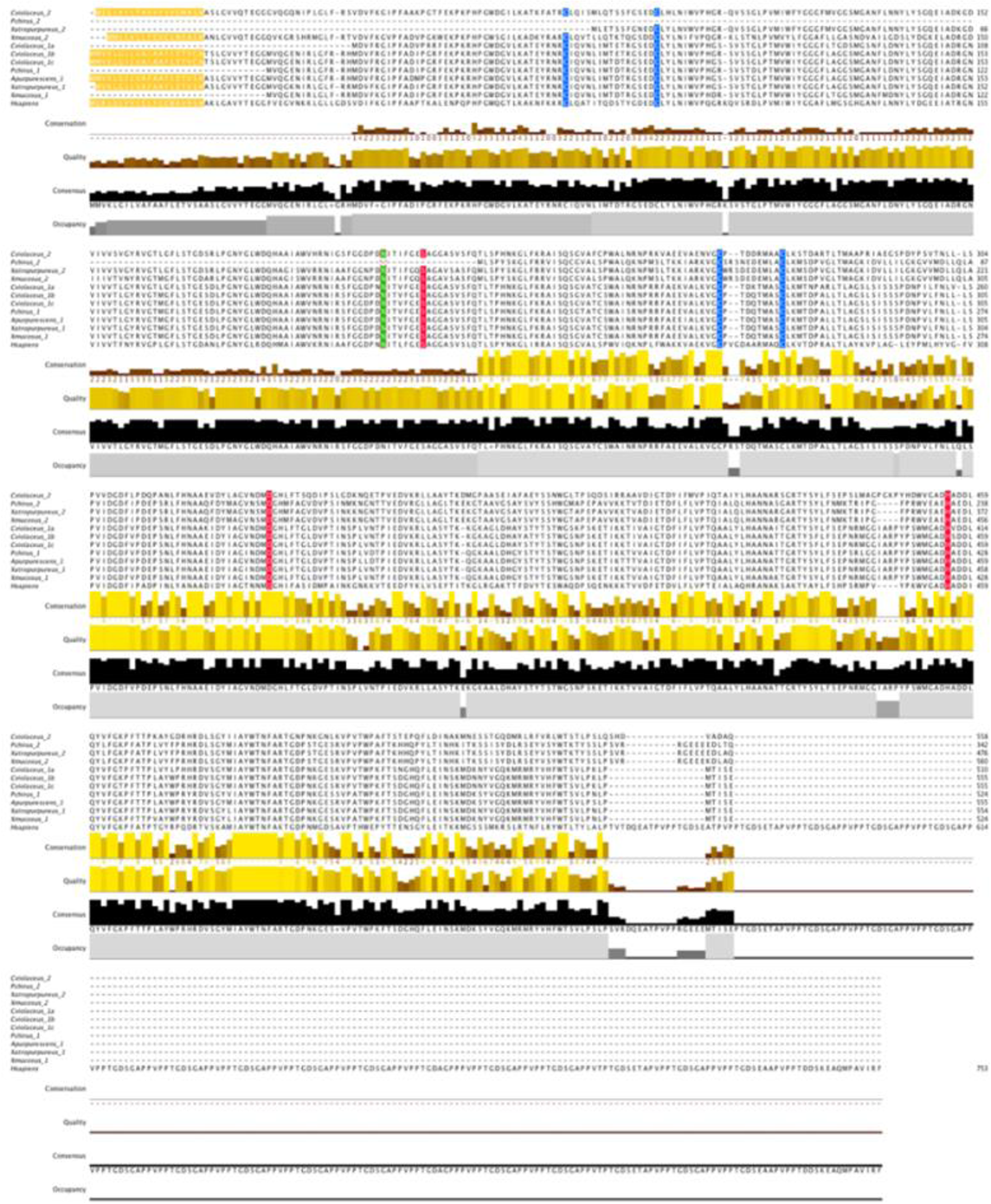
Amino acid sequence alignment of Bile Salt-Activated Lipase (BSAL) sequences. Prickleback and Homo sapiens (as reference) BSAL sequences were aligned in jalview. Amino acids highlighted in yellow represent the signal peptide, blue indicates disulfide bonds, red indicates active sites present, and green indicates glycosylation.

**Supplementary Figure S16:**
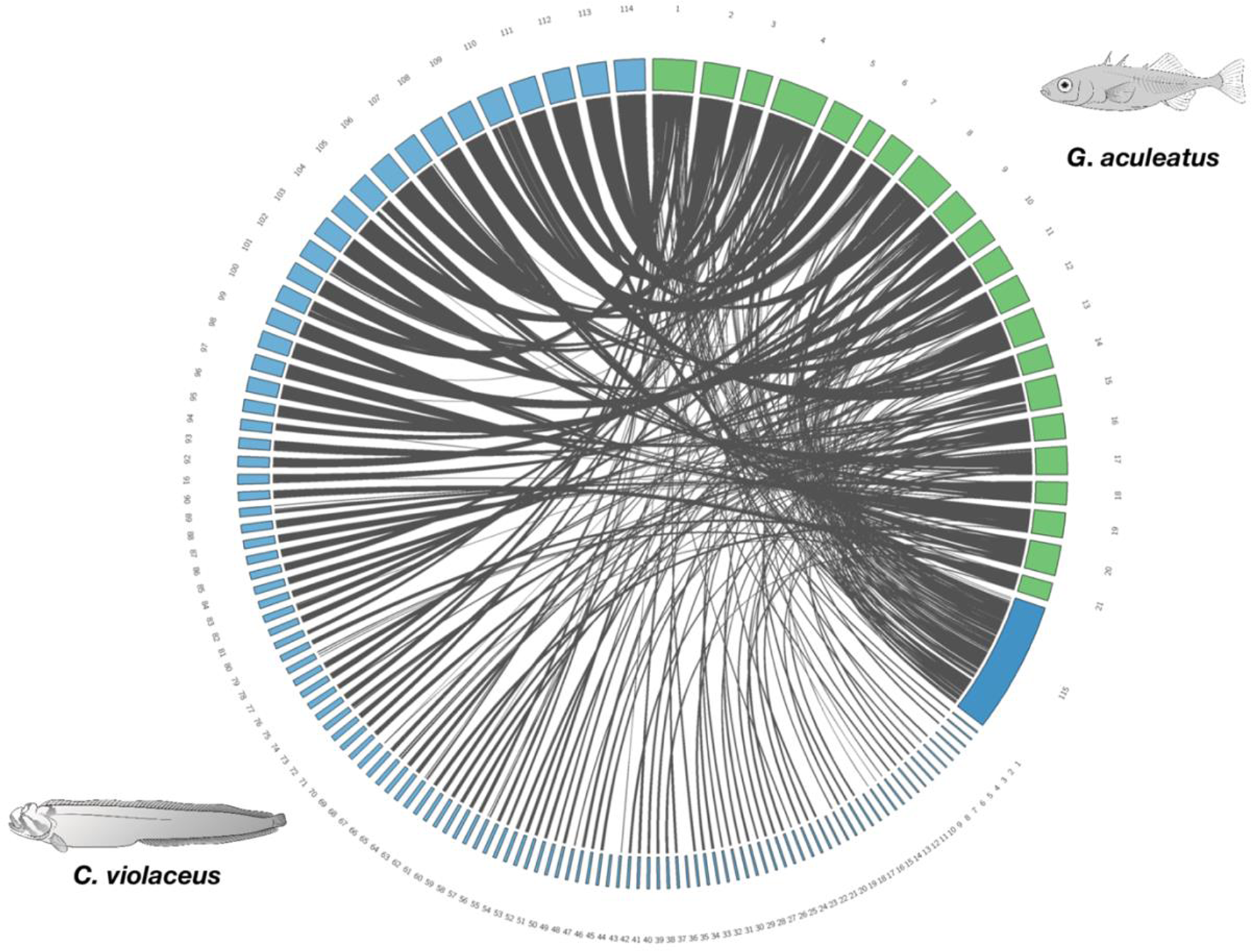
Circos plot showing synteny between the assembled genome *Cebidichthys violaceus* and *Gasterosteus aculeatus* (three-spined stickleback). There are 21 chromosomes (green boxes) which represent the three-spined stickleback genome. There are 114 blue boxes which are 1 MB or greater that represent the *C. violaceus* genome. There are 353 contigs that are less than 1 MB which were concatenated into box labeled as 115. Gray strands indicate syntenic regions between the two genomes. Both *G. aculeatus* and *C. violaceus* illustrations were drawn by Andrea Dingeldein.

**Supplementary Figure S17:**
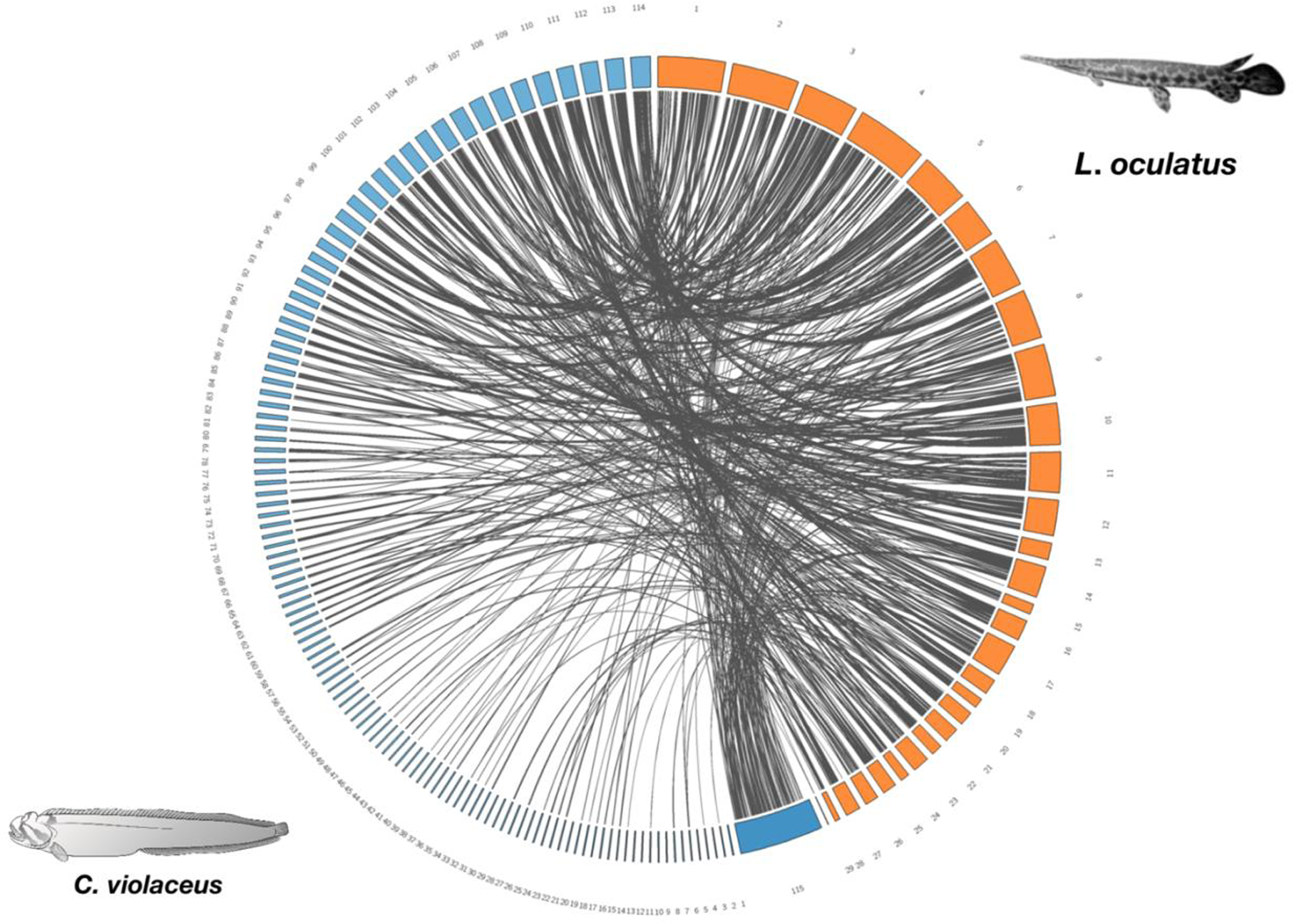
Circos plot showing synteny between the assembled genome *Cebidichthys violaceus* and *Lepisosteus oculatus* (Spotted gar). There are 29 linkage groups (orange boxes) which represent the Spotted gar genome. There are 114 blue boxes which are 1 MB or greater that represent the *C. violaceus* genome. There are 353 contigs that are less than 1 MB which were concatenated into box labeled as 115. Gray strands indicate syntenic regions between the two genomes. *Lepisosteus oculatus* photo was taken by David Solomon and the *C. violaceus* illustration was drawn by Andrea Dingeldein.

**Supplementary Figure S18:**
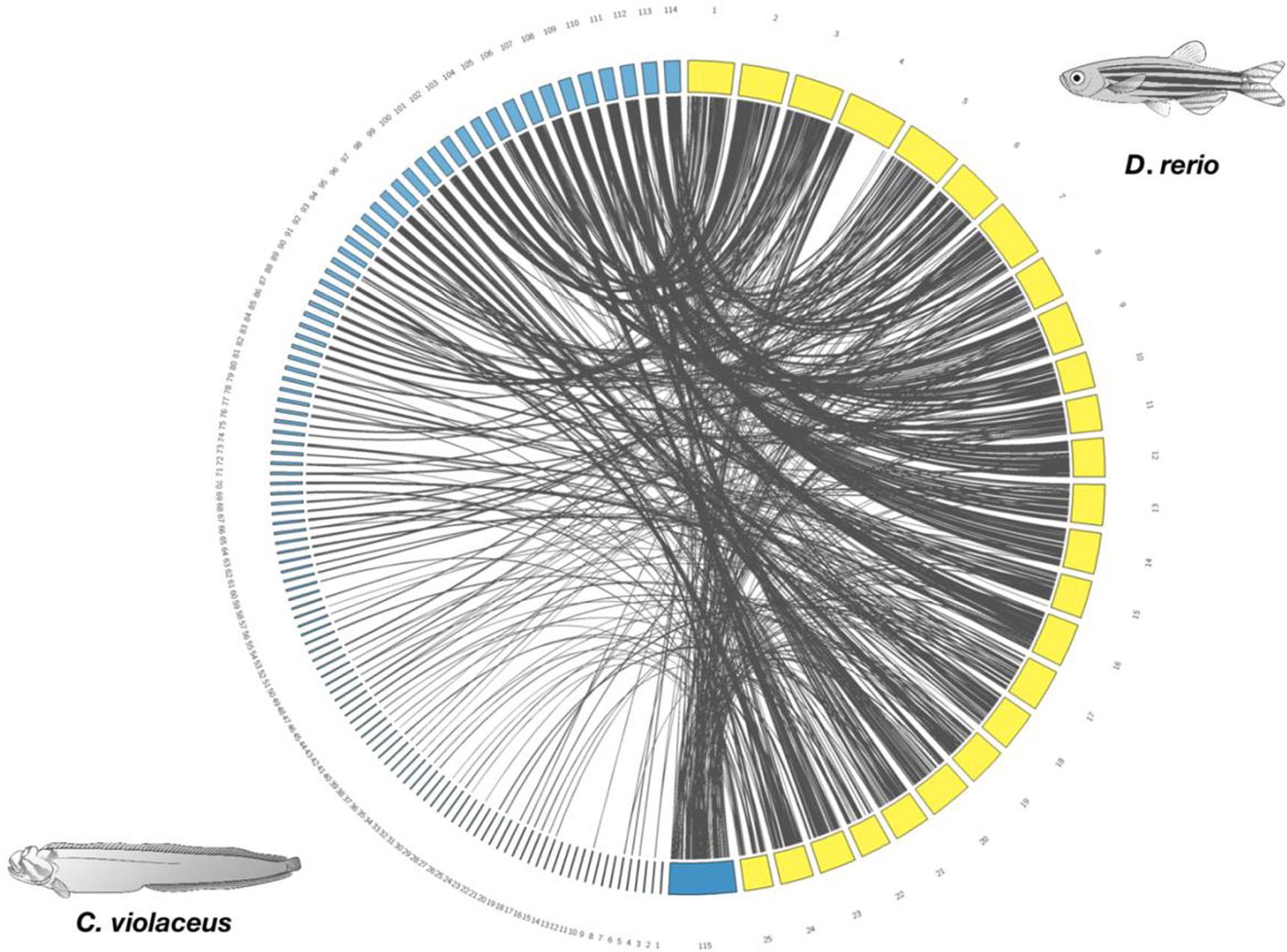
Circos plot showing synteny between the assembled genome *Cebidichthys violaceus* and *Danio rerio* (Zebrafish). There are 25 chromosomes (yellow boxes) which represent the zebrafish genome. There are 114 blue boxes which are 1 MB or greater that represent the *C. violaceus* genome. There are 353 contigs that are less than 1 MB which were concatenated into box labeled as 115. Gray strands indicate syntenic regions between the two genomes. Both *D. rerio* and *C. violaceus* illustrations were drawn by Andrea Dingeldein.

**Supplementary Figure S19:**
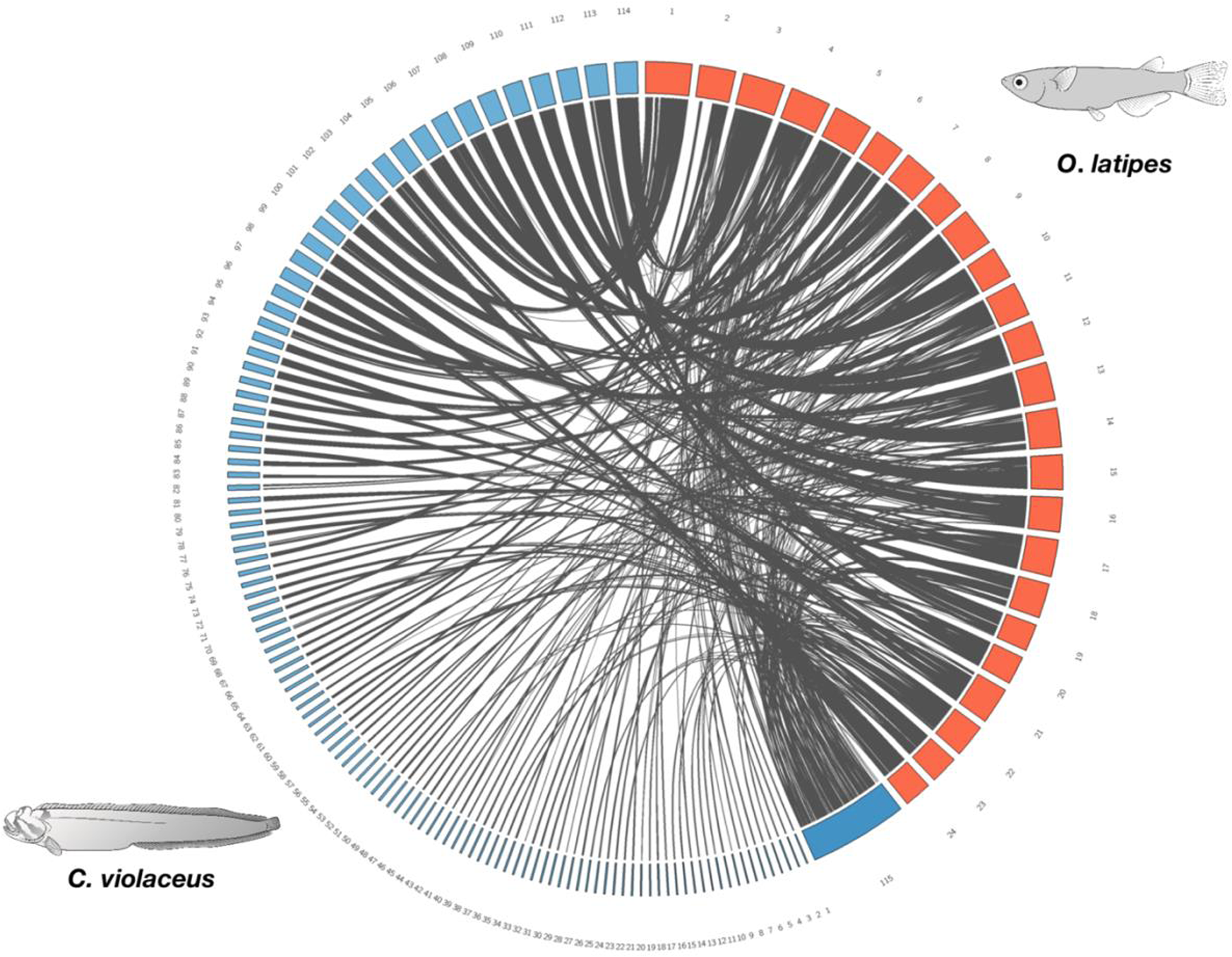
Circos plot showing synteny between the assembled genome *Cebidichthys violaceus* and *Oryzias latipes* (Japanese rice fish). There are 24 chromosomes (red boxes) which represent the Japanese rice fish genome. There are 114 blue boxes which are 1 MB or greater that represent the *C. violaceus* genome. There are 353 contigs that are less than 1 MB which were concatenated into box labeled as 115. Gray strands indicate syntenic regions between the two genomes. Both *O. latipes* and *C. violaceus* illustrations were drawn by Andrea Dingeldein.

**Supplementary Figure S20:**
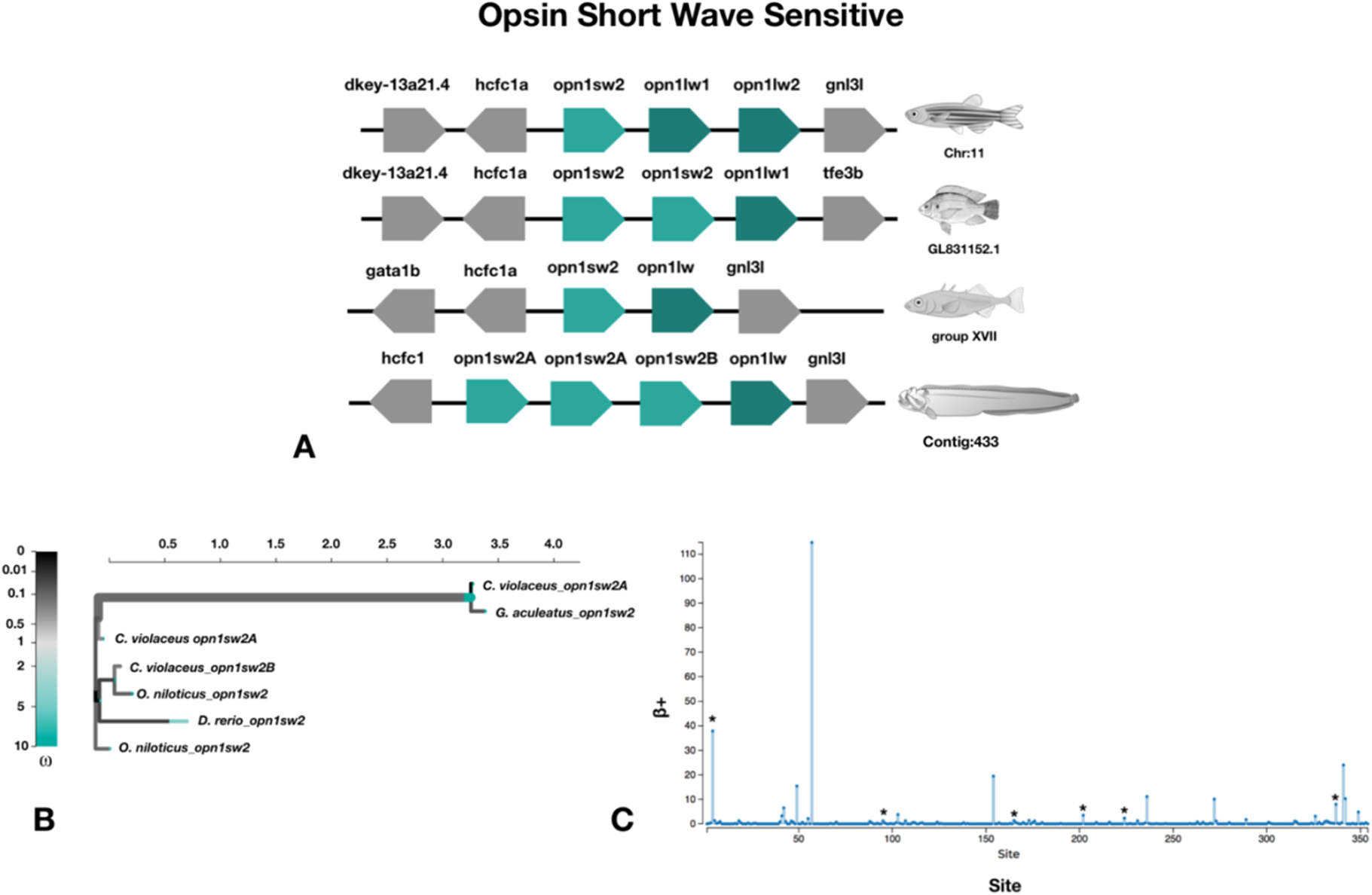
Gene copy number and molecular evolution of Opsin Short Wave Sensitive (opn1sw) genes. a, Synteny map for opn1sw genes from *Danio rerio, Oreochromis niloticus, Gasterosteus aculeatus*, and *Cebidichthys violaceus. D. rerio, G. aculeatus*, and *C. violaceus* were drawn by Andrea Dingeldein. *O. niloticus* illustration was was taken from www.fao.org. b, An adaptive Branch-Site Random Effects Likelihood (aBSREL) test for episodic diversification was estimated and represented as a phylogenetic tree for opn1sw genes from *C. violaceus* and three other fishes. ω is the ratio of nonsynonymous to synonymous substitutions. The color gradient represents the magnitude of the corresponding ω. Branches thicker than the other branches have a P<0.05 (corrected for multiple testing) to reject the null hypothesis of all ω on that branch (neutral or negative selection only). A thick branch is considered to have experienced diversifying positive selection. c, The output of Mixed Efffects Model of Evolution (MEME) to detect episodic positive/diversifying selection at sites. β+ is the non-synonymous substitution rate at a site for the positive/neutral evolution throughout the sequence of the gene. ** is an indication that the positive/diversifying site is statistically significant with a p-value < 0.01 and * is for p-value < 0.05.

**Supplementary Table S2:**
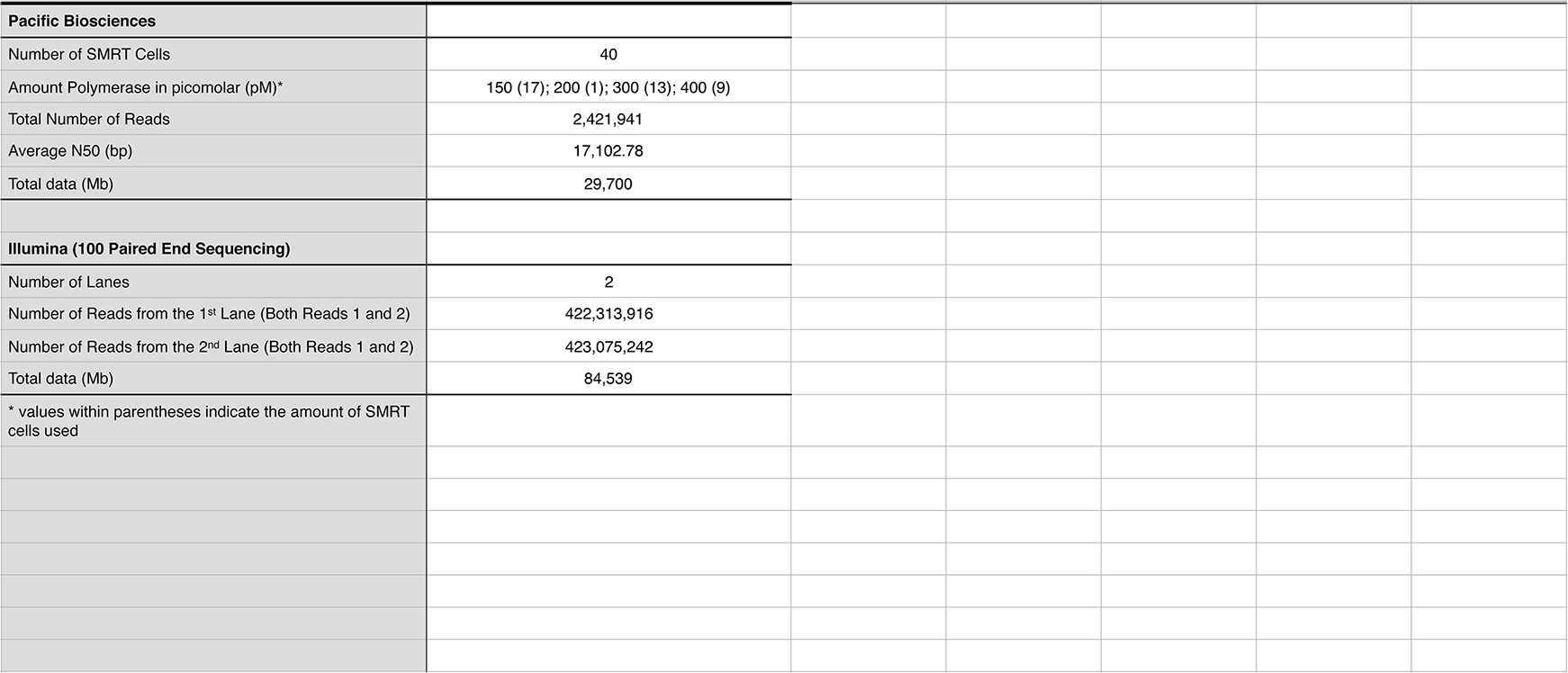
Genome Sequencing Information

**Supplementary Table S3:**
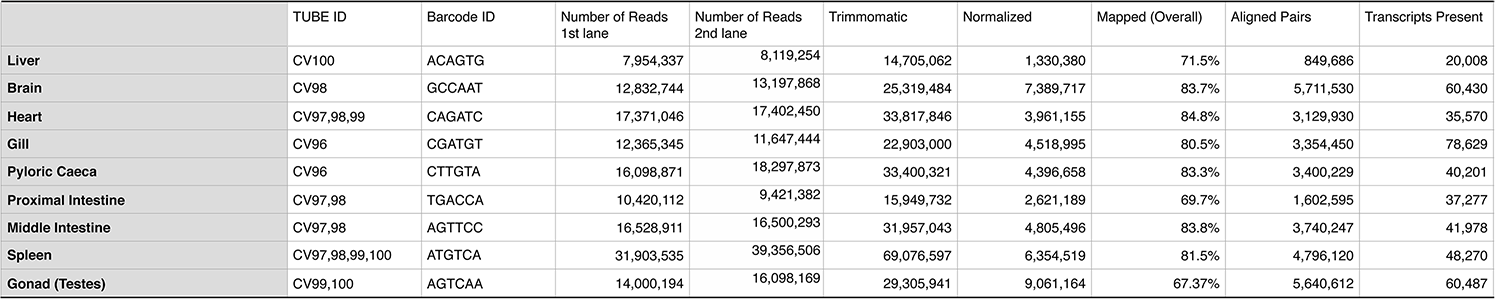
Transcriptomic Sequencing Information and Trinity Assembly

**Supplementary Table S4:** RepeatMasker for the *C. violaceus* assembled genome

|  |  |  |  |
| --- | --- | --- | --- |
|  | Number of Elements | Length Occupied (bp) | Percentage of Sequence (%) |
| Retroelements | 11715 | 5340689 | 0.9 |
| SINEs: | 1985 | 223655 | 0.04 |
| Penelope | 60 | 24362 | 0.0 |
| LINEs: | 8905 | 4191047 | 0.71 |
| CRE/SLACS | 0 | 0 | 0.0 |
| L2/CR1/Rex | 4980 | 2049678 | 0.35 |
| R1/LOA/Jockey | 0 | 0 | 0.0 |
| R2/R4/NeSL | 85 | 24828 | 0.0 |
| RTE/Bov-B | 3316 | 1768649 | 0.3 |
| L1/CIN4 | 336 | 205561 | 0.03 |
| LTR elements: | 825 | 925987 | 0.16 |
| BEL/Pao | 20 | 36895 | 0.01 |
| Ty1/Copia | 22 | 20106 | 0.0 |
| Gypsy/DIRS1 | 703 | 846315 | 0.14 |
| Retroviral | 79 | 22637 | 0.0 |
| DNA transposons | 10562 | 2582768 | 0.44 |
| hobo-Activator | 4384 | 730619 | 0.12 |
| Tc1-IS630-Pogo | 4812 | 1690957 | 0.29 |
| En-Spm | 0 | 0 | 0.0 |
| MuDR-IS905 | 0 | 0 | 0.0 |
| PiggyBac | 352 | 30854 | 0.01 |
| Tourist/Harbinger | 178 | 26872 | 0.00 |
| Other (Mirage, P-element, Transib) | 0 | 0 | 0.00 |
| Rolling-circles | 0 | 0 | 0.0 |
| Unclassified: | 203 | 21708 | 0.0 |
| Total interspersed repeats: |  | 7945165 | 1.34 |
| Small RNA: | 2430 | 215618 | 0.04 |
| Satellites: | 4 | 1112 | 0.00 |
| Simple repeats: | 468887 | 27486227 | 4.64 |
| Low complexity: | 37947 | 2434577 | 0.41 |

**Supplementary Table S5:**
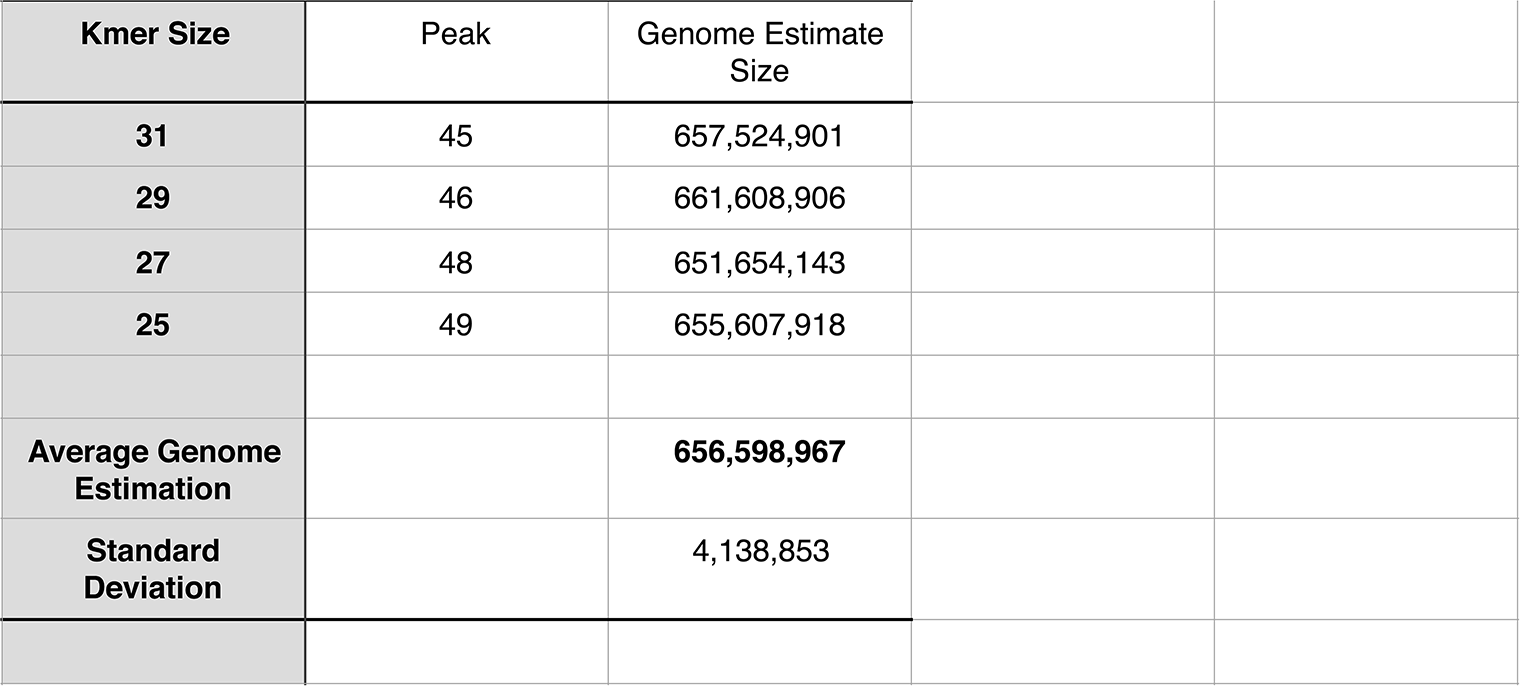
Genome Size Estimation with Jellyfish v2.2.0 (Marçais and Kingsford, 2011)

**Supplementary Table S6:**
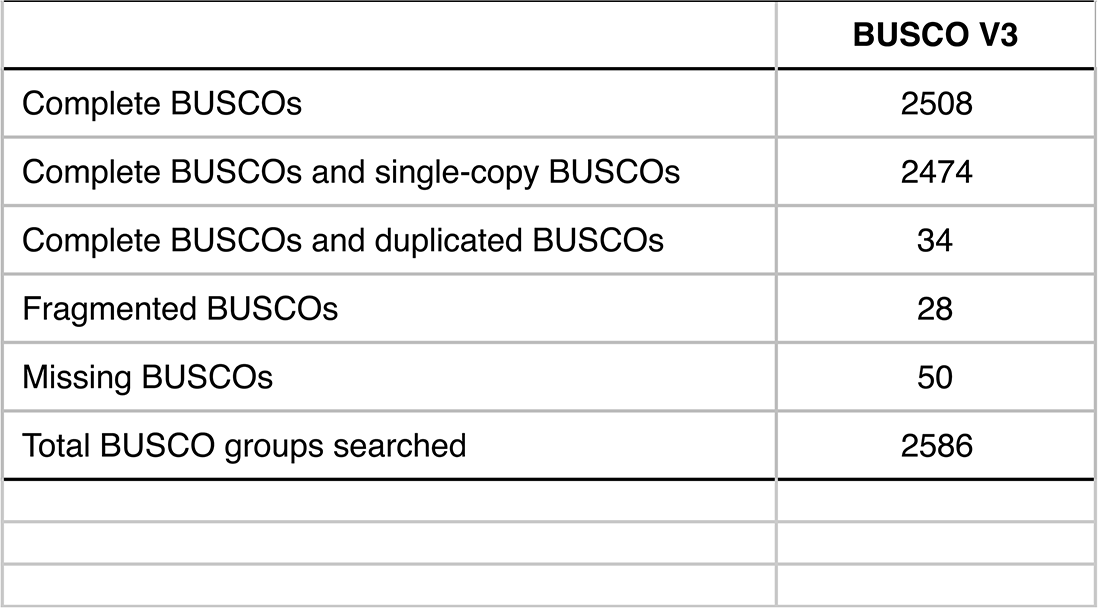
BUSCO v3 Estimation on the *Cebidichthys violaceus* genome

**Supplementary Table S7:**
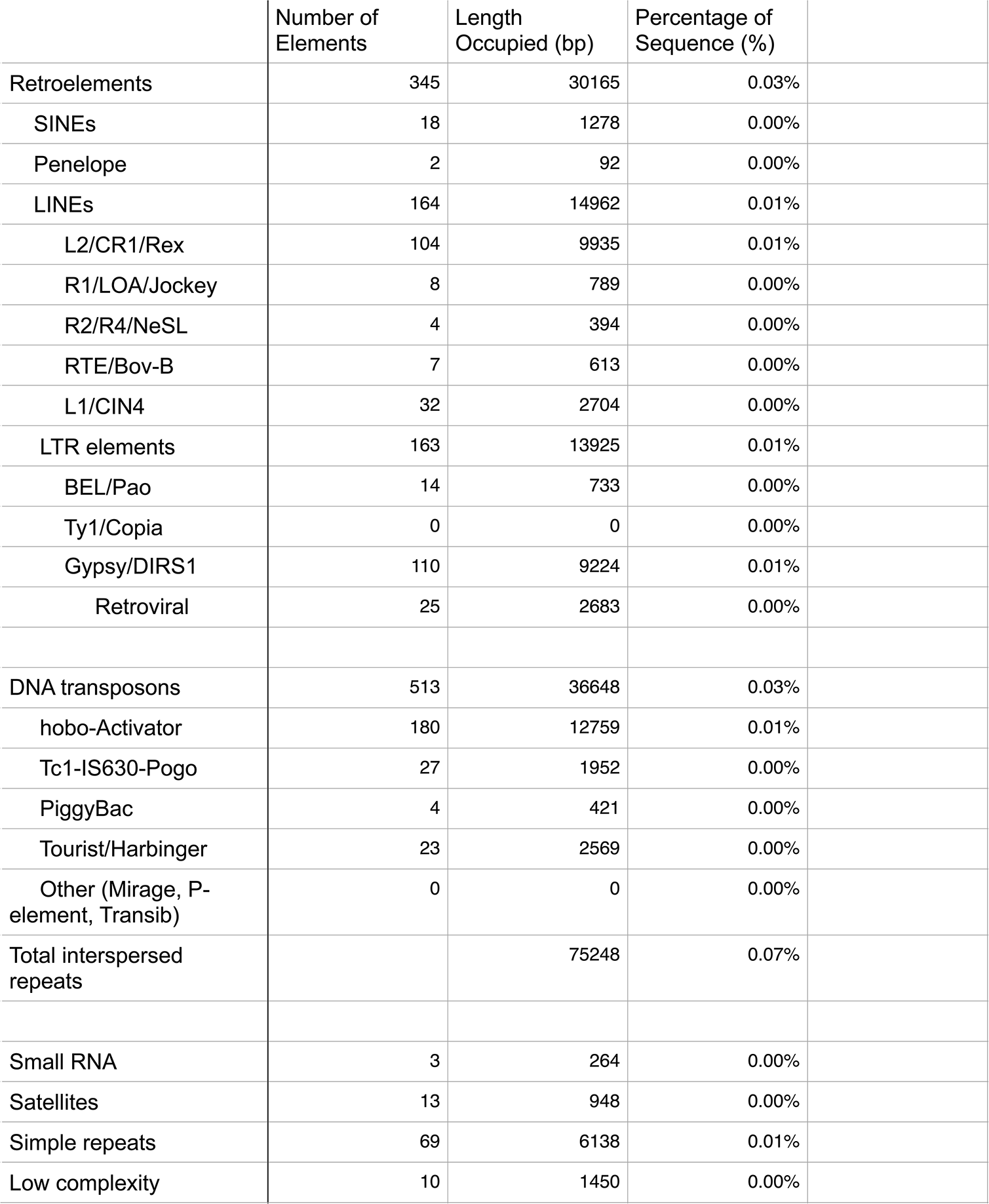
RepeatMasker for the Genome Guided (TRINITY) Transcriptome

**Supplementary Table S8:**
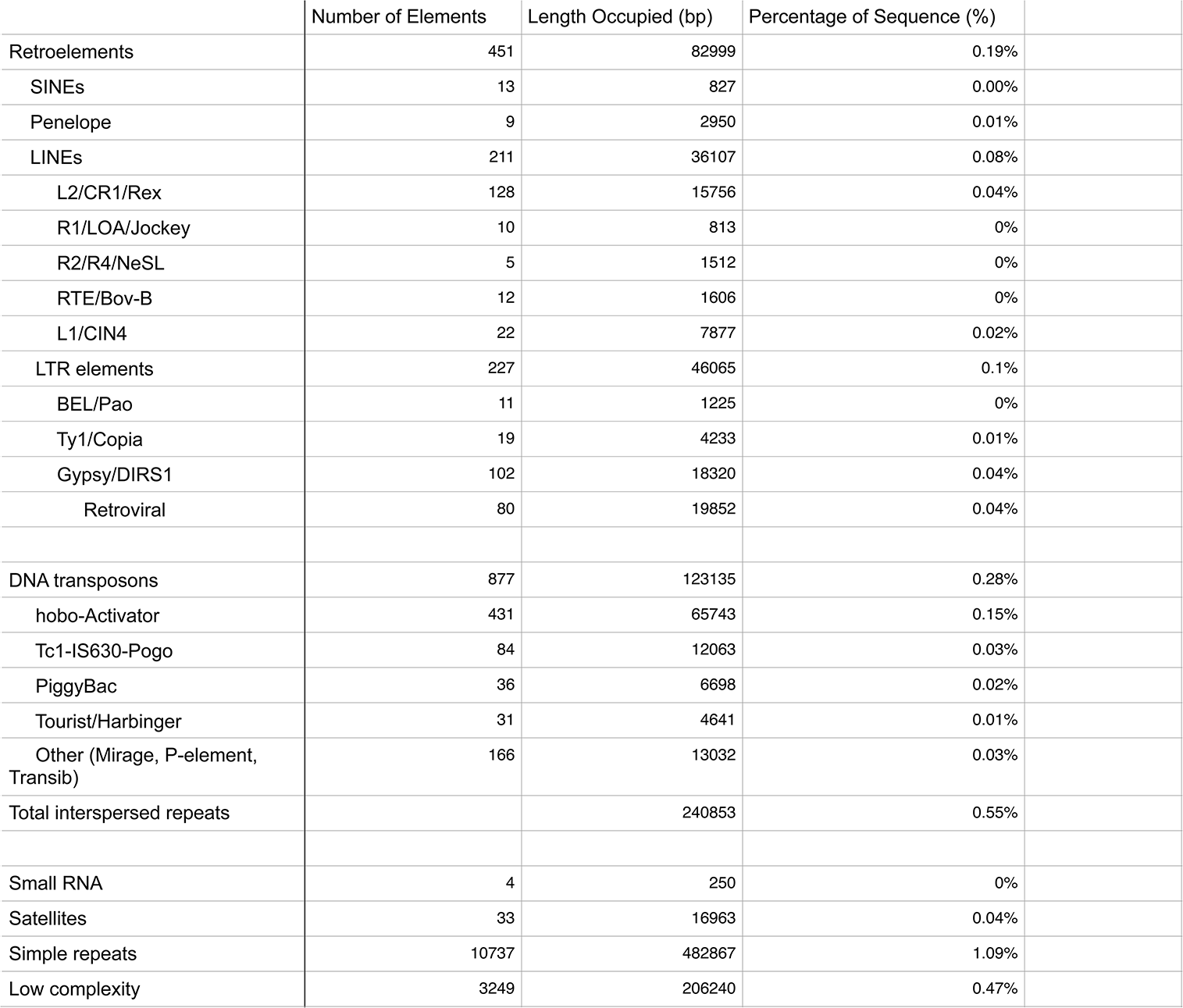
RepeatMasker for Augustus Predicted Genes

**Supplementary Table S9:** Estimation of Full Length of Transcripts from all nine transcriptomes

| **Hit percent coverage bin** | **Count in bin** | **>Bin below** |
| --- | --- | --- |
| 100 | 2692 | 2692 |
| 90 | 1258 | 3950 |
| 80 | 1249 | 5199 |
| 70 | 1358 | 6557 |
| 60 | 1598 | 8155 |
| 50 | 1991 | 10146 |
| 40 | 2411 | 12557 |
| 30 | 2912 | 15469 |
| 20 | 2966 | 18435 |
| 10 | 1132 | 19567 |

**Supplementary Table S10:**
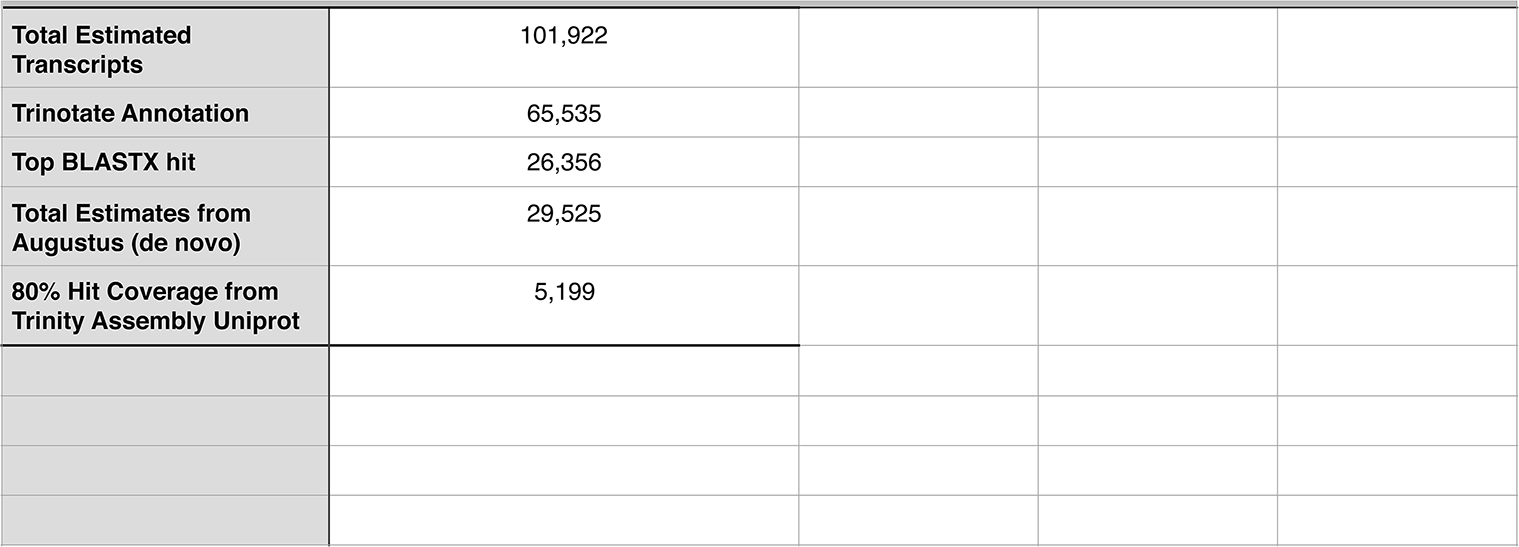
Estimation of Total Genes from Trinity and Augustus

**Supplementary Table S11:**
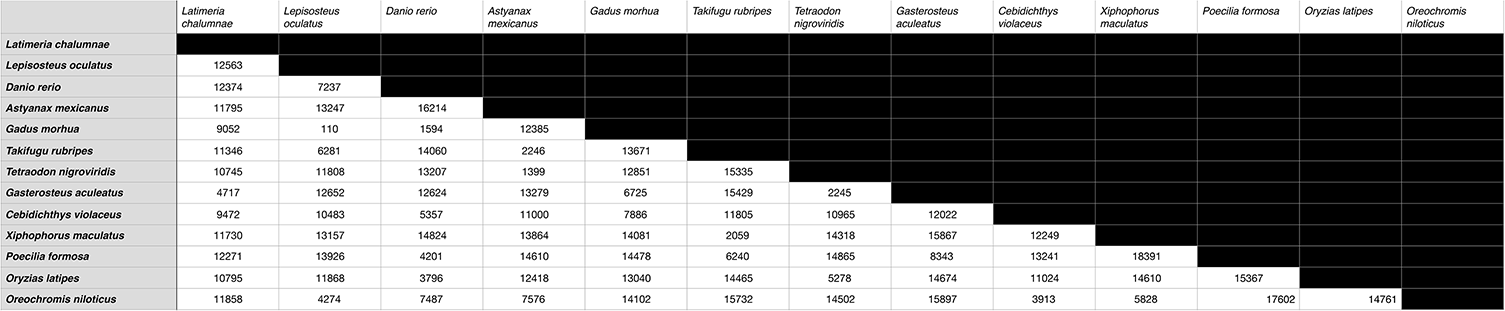
Pairwise Comparison of Orthologs of Cebidichthys violaceus and fish genomes deposited on Ensembl

## ACKNOWLEDGEMENTS

We would like to thank H. Yip and Q.B. Nguyen-Phuc for assistance in collected samples for genomic and transcriptomic analyses. D. Canestro for providing assistance and facilities at the Kenneth S. Norris Rancho Marino Reserve in Cambria, CA while collecting samples for this project. We would like to thank M. Oakes, V. Ciobanu, S.A. Chung, D. Yu, and Y. Kanomata at the UC Irvine Genomics High-Throughput Facility. A. Long, J. Baldwin-Brown, and K. Thorton for RNA-Seq assembly suggestions. The comparative physiology group at UC Irvine for providing feedback on the content of this manuscript. We would also like to thank N. Nirale for assistance in uploading all genomic and transcriptomic sequences onto NCBI’s GenBank. All sequence data was deposited to NCBI’s GenBank under the bioproject ID: PRJNA384078. Included under this bioproject are the Illumina genomic and tissue transcriptomic objects under SAMN06857690-SAMN0687699, 40 SMRT cells of PacBio sequencing under SAMN06857690, and the final genome assembly (NJBE00000000). This project was funded through NSF-IOS 1355224. Lastly, authors declare no conflicts of interests.

## Bibliography

Altschul, S. Gapped BLAST and PSI-BLAST: a new generation of protein database search programs. Nucleic Acids Research 25, 3389–3402 (1997).

Ashburner, M. et al. Gene Ontology: tool for the unification of biology. Nature genetics 25, 25 (2000).

Axelsson, E. et al. The genomic signature of dog domestication reveals adaptation to a starch-rich diet. Nature 495, 360–364 (2013).

Benson, G. Tandem repeats finder: a program to analyze DNA sequences. Nucleic Acids Research 27, 573–580 (1999).

Bergman, E. N. Energy contributions of volatile fatty acids from the gastrointestinal tract in various species. Physiological Reviews 70, 567–590 (1990).

Boeckmann, B. The SWISS-PROT protein knowledgebase and its supplement TrEMBL in 2003. Nucleic Acids Research 31, 365–370 (2003).

Boehlke, C., Zierau, O. & Hannig, C. Salivary amylase – The enzyme of unspecialized euryphagous animals. Archives of Oral Biology 60, 1162–1176 (2015).

Bolger, A. M., Lohse, M. & Usadel, B. Trimmomatic: a flexible trimmer for Illumina sequence data. Bioinformatics 30, 2114–2120 (2014).

Buddington, R. K. & Diamond, J. M. Pyloric ceca of fish: a “new” absorptive organ. American Journal of Physiology-Gastrointestinal and Liver Physiology 252(1), G65–G76. (1987).

Buddington, R. K., Chen, J. W. & Diamond, J. Genetic and phenotypic adaptation of intestinal nutrient transport to diet in fish. The Journal of Physiology 393, 261–281 (1987).

Cai, Wenjing, et al. Different strategies of grass carp *(Ctenopharyngodon idella)* responding to insufficient or excessive dietary carbohydrate. Aquaculture 497, 292–298 (2018).

Chakraborty, M. & Fry, J. D. Parallel Functional Changes in Independent Testis-Specific Duplicates of Aldehyde dehydrogenase in *Drosophila*. Molecular Biology and Evolution 32, 1029–1038 (2015).

Chakraborty, M., Baldwin-Brown, J. G., Long, A. D. & Emerson, J. J. Contiguous and accurate de novo assembly of metazoan genomes with modest long read coverage. Nucleic Acids Research 44, e147-e147 (2016). doi:10.1093/nar/gkw654

Chan, A. S., Horn, M. H., Dickson, K. A. & Gawlicka, A. Digestive enzyme activities in carnivores and herbivores: comparisons among four closely related prickleback fishes (Teleostei: Stichaeidae) from a California rocky intertidal habitat. Journal of Fish Biology 65, 848–858 (2004).

Chen, Z., Farrell, A. P., Matala, A. & Narum, S. R. Mechanisms of thermal adaptation and evolutionary potential of conspecific populations to changing environments. Molecular Ecology 27, 659–674 (2018).

Chin, C. S. et al. Nonhybrid, finished microbial genome assemblies from long-read SMRT sequencing data. Nature methods 10, 563 (2013).

Choat, J. H. & Clements, K. D. Vertebrates herbivores in marine and terrestrial environments: A Nutritional Ecology Perspective. Annual Review of Ecology and Systematics 29, 375–403 (1998).

Clements, K. D., & Raubenheimer, D. (2006). Feeding and nutrition. The Physiology of fishes, 47–82.

Clements, K. D., German, D. P., Piché, J., Tribollet, A. & Choat, J. H. Integrating ecological roles and trophic diversification on coral reefs: multiple lines of evidence identify parrotfishes as microphages. Biological Journal of the Linnean Society 120, 729–751. (2017). doi:10.1111/bij.12914

Clissold, F. J., Tedder, B. J., Conigrave, A. D. & Simpson, S. J. The gastrointestinal tract as a nutrient-balancing organ. Proceedings of the Royal Society B: Biological Sciences 277, 1751–1759 (2010).

Dasmahapatra, K. K., Walters, J. R., Briscoe, A. D., Davey, J. W., Whibley, A., Nadeau, N. J., … & Salazar, C. Butterfly genome reveals promiscuous exchange of mimicry adaptations among species. Nature 487(7405), 94 (2012).

Edgar, R. C. MUSCLE: a multiple sequence alignment method with reduced time and space complexity. BMC bioinformatics 5, 113 (2004).

Feschotte, C. & Pritham, E. J. DNA Transposons and the Evolution of Eukaryotic Genomes. Annual Review of Genetics 41, 331–368 (2007).

Finn, R. D. et al. The Pfam protein families database. Nucleic Acids Research 36, (2007).

German, D. P., Foti, D. M., Heras, J., Amerkhanian, H. & Lockwood, B. L. Elevated Gene Copy Number Does Not Always Explain Elevated Amylase Activities in Fishes. Physiological and Biochemical Zoology 89, 277–293 (2016).

German, D. P., Horn, M. H., & Gawlicka, A. Digestive enzyme activities in herbivorous and carnivorous prickleback fishes (Teleostei: Stichaeidae): ontogenetic, dietary, and phylogenetic effects. Physiological and Biochemical Zoology 77, 789–804 (2004).

German, D. P., Neuberger, D. T., Callahan, M. N., Lizardo, N. R. & Evans, D. H. Feast to famine: The effects of food quality and quantity on the gut structure and function of a detritivorous catfish (Teleostei: Loricariidae). Comparative Biochemistry and Physiology Part A: Molecular & Integrative Physiology 155, 281–293 (2010).

German, D. P., Sung, A., Jhaveri, P. & Agnihotri, R. More than one way to be an herbivore: convergent evolution of herbivory using different digestive strategies in prickleback fishes (Stichaeidae). Zoology 118, 161–170 (2015).

German, J. et al. Effect of dietary fats and barley fiber on total cholesterol and lipoprotein cholesterol distribution in plasma of hamsters. Nutrition Research 16, 1239–1249 (1996).

Gioda, C. R. et al. Different feeding habits influence the activity of digestive enzymes in freshwater fish. Ciência Rural 47, (2017).

Grabherr, M. G. et al. Full-length transcriptome assembly from RNA-Seq data without a reference genome. Nature Biotechnology 29, 644–652 (2011).

Grabherr, M. G. et al. Genome-wide synteny through highly sensitive sequence alignment: Satsuma. Bioinformatics 26, 1145–1151 (2010).

Guindon, S. et al. New Algorithms and Methods to Estimate Maximum-Likelihood Phylogenies: Assessing the Performance of PhyML 3.0. Systematic Biology 59, 307–321 (2010).

Harris, S. E. & Munshi-South, J. Signatures of positive selection and local adaptation to urbanization in white-footed mice (Peromyscus leucopus). Molecular Ecology 26, 6336–6350 (2016). doi: 10.1101/038141

Herrel, A. et al. Rapid large-scale evolutionary divergence in morphology and performance associated with exploitation of a different dietary resource. Proceedings of the National Academy of Sciences 105, 4792–4795 (2008).

Hinegardner, R., & Rosen, D. E. Cellular DNA content and the evolution of teleostean fishes. The American Naturalist 106, 621–644 (1972).

Horn, M. H. et al. Structure and function of the stomachless digestive system in three related species of New World silverside fishes (Atherinopsidae) representing herbivory, omnivory, and carnivory. Marine Biology 149, 1237–1245 (2006).

Horn, M. H., & Riegle, K. C. Evaporative water loss and intertidal vertical distribution in relation to body size and morphology of stichaeoid fishes from California. Journal of Experimental Marine Biology and Ecology 50, 273–288. (1981).

Horn, M. H., Neighbors, M. A. & Murray, S. N. Herbivore responses to a seasonally fluctuating food supply: Growth potential of two temperate intertidal fishes based on the protein and energy assimilated from their macroalgal diets. Journal of Experimental Marine Biology and Ecology 103, 217–234 (1986).

Hsieh, P. et al. Exome Sequencing Provides Evidence of Polygenic Adaptation to a Fat-Rich Animal Diet in Indigenous Siberian Populations. Molecular Biology and Evolution 34, 2913–2926 (2017).

Kajitani, R. et al. Efficient de novo assembly of highly heterozygous genomes from whole-genome shotgun short reads. Genome Research 24, 1384–1395 (2014).

Karasov, W. H., & del Rio, C. M. Physiological ecology: how animals process energy, nutrients, and toxins. Princeton University Press. (2007).

Kim, D. et al. TopHat2: accurate alignment of transcriptomes in the presence of insertions, deletions and gene fusions. Genome Biology 14, R36 (2013).

Kim, Ji Hyun, et al. Alginate/bacterial cellulose nanocomposite beads prepared using Gluconacetobacter xylinus and their application in lipase immobilization. Carbohydrate polymers 157, 137–145 (2017).

Kim, K. H., Horn, M. H., Sosa, A. E. & German, D. P. Sequence and expression of an α-amylase gene in four related species of prickleback fishes (Teleostei: Stichaeidae): ontogenetic, dietary, and species-level effects. Journal of Comparative Physiology B 184, 221–234 (2014).

Kohl, K. D., Weiss, R. B., Dale, C. & Dearing, M. D. Diversity and novelty of the gut microbial community of an herbivorous rodent (Neotoma bryanti). Symbiosis 54, 47–54 (2011).

Krzywinski, M. et al. Circos: An information aesthetic for comparative genomics. Genome Research 19, 1639–1645 (2009).

Kumar, S., Stecher, G., & Tamura, K. MEGA7: molecular evolutionary genetics analysis version 7.0 for bigger datasets. Molecular biology and evolution 33, 1870–1874 (2016).

Kurtz, S. et al. Versatile and open software for comparing large genomes. Genome Biology 5, R12 (2004).

Lagesen, K. et al. RNAmmer: consistent and rapid annotation of ribosomal RNA genes. Nucleic Acids Research 35, 3100–3108 (2007).

Lamichhaney, S. et al. Evolution of Darwin’s finches and their beaks revealed by genome sequencing. Nature 518, 371–375 (2015).

Langmead, B. & Salzberg, S. L. Fast gapped-read alignment with Bowtie 2. Nature Methods 9, 357–359 (2012).

Lawrence, T. J. et al. FAST: FAST Analysis of Sequences Toolbox. Frontiers in Genetics 6, (2015).

Leigh, S. C., Papastamatiou, Y. P., & German, D. P. Seagrass digestion by a notorious ‘carnivore’. Proceedings of the Royal Society B 285, 20181583. (2018).

Li, H. et al. The Sequence Alignment/Map format and SAMtools. Bioinformatics 25, 2078–2079 (2009).

Marçais, G. & Kingsford, C. A fast, lock-free approach for efficient parallel counting of occurrences of k-mers. Bioinformatics 27, 764–770 (2011).

Mcvean, G. What drives recombination hotspots to repeat DNA in humans? Philosophical Transactions of the Royal Society B: Biological Sciences 365, 1213–1218 (2010).

Mendoza, M. L. Z., et al. Hologenomic adaptations underlying the evolution of sanguivory in the common vampire bat. Nature ecology & evolution 2, 659 (2018).

Min, X. J., Butler, G., Storms, R. & Tsang, A. OrfPredictor: predicting protein-coding regions in EST-derived sequences. Nucleic Acids Research 33, (2005).

Murray, H. M., Gallant, J. W., Perez-Casanova, J. C., Johnson, S. C. & Douglas, S. E. Ontogeny of lipase expression in winter flounder. Journal of Fish Biology 62, 816–833 (2003).

Neighbors, M. A. & Horn, M. H. Nutritional quality of macrophytes eaten and not eaten by two temperatezone herbivorous fishes: A multivariate comparison. Marine Biology 108, 471–476 (1991).

Obrien, K. P. Inparanoid: a comprehensive database of eukaryotic orthologs. Nucleic Acids Research 33, (2005).

Painter, T. J. Algal Polysaccharides. The Polysaccharides 195–285 (1983). doi:10.1016/b978-0-12-065602-8.50009-1

Pantzartzi, C. N., Pergner, J. & Kozmik, Z. The role of transposable elements in functional evolution of amphioxus genome: the case of opsin gene family. Scientific Reports 8, (2018).

Peichel, C. L. & Marques, D. A. The genetic and molecular architecture of phenotypic diversity in sticklebacks. Philosophical Transactions of the Royal Society B: Biological Sciences 372, 20150486 (2016).

Perry, G. H. et al. Diet and the evolution of human amylase gene copy number variation. Nature Genetics 39, 1256–1260 (2007).

Petersen, T. N., Brunak, S., Heijne, G. V. & Nielsen, H. SignalP 4.0: discriminating signal peptides from transmembrane regions. Nature Methods 8, 785–786 (2011).

Powell, S. et al. eggNOG v4.0: nested orthology inference across 3686 organisms. Nucleic Acids Research 42, (2013).

Protas, M. E., et al. Genetic analysis of cavefish reveals molecular convergence in the evolution of albinism. Nature genetics 38, 107 (2006).

Rungruangsak-Torrissen, K., Moss, R., Andresen, L. H., Berg, A., & Waagbø, R. Different expressions of trypsin and chymotrypsin in relation to growth in Atlantic salmon (Salmo salar L.). Fish Physiology and Biochemistry 32, 7 (2006).

Simão, F. A., Waterhouse, R. M., Ioannidis, P., Kriventseva, E. V., & Zdobnov, E. M. BUSCO: assessing genome assembly and annotation completeness with single-copy orthologs. Bioinformatics 31, 3210–3212 (2015).

Smit, A. F. Repeat-Masker Open-3.0. http://www.repeatmasker.org. (2004).

Stanke, M. & Morgenstern, B. AUGUSTUS: a web server for gene prediction in eukaryotes that allows user-defined constraints. Nucleic Acids Research 33, (2005).

Stanke, M., Keller, O., Gunduz, I., Hayes, A., Waack, S., Morgenstern, B. AUGUSTUS: ab initio prediction of alternative transcripts. Nucleic Acids Research 34, W435–W439 (2006).

Tengjaroenkul, B., Smith, B. J., Caceci, T. & Smith, S. A. Distribution of intestinal enzyme activities along the intestinal tract of cultured Nile tilapia, Oreochromis niloticus L. Aquaculture 182, 317–327 (2000).

Tong, C., Tian, F. & Zhao, K. Genomic signature of highland adaptation in fish: a case study in Tibetan Schizothoracinae species. BMC Genomics 18, (2017).

Trapnell, C. et al. Differential gene and transcript expression analysis of RNA-seq experiments with TopHat and Cufflinks. Nature Protocols 7, 562–578 (2012).

Vega-Retter, C. et al. Differential gene expression revealed with RNA-Seq and parallel genotype selection of the ornithine decarboxylase gene in fish inhabiting polluted areas. Scientific Reports 8, (2018).

Walker, B. J. et al. Pilon: An Integrated Tool for Comprehensive Microbial Variant Detection and Genome Assembly Improvement. PLoS ONE 9, (2014).

Waterhouse, R. M. et al. BUSCO Applications from Quality Assessments to Gene Prediction and Phylogenomics. Molecular Biology and Evolution 35, 543–548 (2017).

Weaver, S. et al. Datamonkey 2.0: A Modern Web Application for Characterizing Selective and Other Evolutionary Processes. Molecular Biology and Evolution 35, 773–777 (2018).

Willmott, M. E., Clements, K. D. & Wells, R. M. The influence of diet and gastrointestinal fermentation on key enzymes of substrate utilization in marine teleost fishes. Journal of Experimental Marine Biology and Ecology 317, 97–108 (2005).

Ye, C., Hill, C. M., Wu, S., Ruan, J. & Ma, Z. DBG2OLC: Efficient Assembly of Large Genomes Using Long Erroneous Reads of the Third Generation Sequencing Technologies. Scientific Reports 6, 31900 (2016).

Zerbino, D. R. et al. Ensembl 2018. Nucleic acids research 46, D754–D761 (2017).

